# A neural circuit framework for economic choice: from building blocks of valuation to compositionality in multitasking

**DOI:** 10.1101/2025.03.13.643098

**Authors:** Aldo Battista, Camillo Padoa-Schioppa, Xiao-Jing Wang

## Abstract

Value-guided decisions are a cornerstone of cognition, yet the underlying circuit-level mechanisms remain elusive. We used reinforcement learning to train recurrent neural network models endowed with Dale’s law on a battery of economic choice tasks, which revealed a two-stage computational framework. First, value estimation occurs at input level where learned weights store subjective preferences and approximate the nonlinear multiplication of reward magnitude and probability to yield expected values. This feedforward mechanism enables generalization to novel choice options. Second, option values are compared within the recurrent network, where specific connectivity patterns mediate robust winner-take-all decisions, with both excitatory and inhibitory neurons exhibiting value and choice selectivity. By training a single network on multiple tasks, we show compositional representations combining a shared computational schema with specialized neural modules. Reproducing key neurophysiological findings from the primate orbitofrontal cortex, our model unifies value computation, comparison, and generalization into a coherent framework with testable predictions.

**IN BRIEF:** Battista et al. use biologically plausible RNNs to uncover circuit mechanisms of economic choice. They propose a two-stage framework where feedforward inputs compute offer values and recurrent inhibition drives comparison. This architecture explains how the brain generalizes preferences and multitasks using compositional neural codes, offering a unified theory of decisionmaking.

**HIGHLIGHTS:**

- Biologically plausible RNN reveals a two-stage economic choice framework
- Input weights store preferences and multiply features to compute offer value
- Competitive recurrent inhibition mediates winner-take-all comparison
- Multitasking relies on compositional shared and specialized neural modules

## INTRODUCTION

Economic choice—making decisions based on subjective preferences—is fundamental to human and animal behavior^1,2^.

Neuroscientific studies of economic choice at the single-cell level began around the turn of this century^3–6^. Understanding economic decisions relies on *subjective value*, a metric facilitating comparison between different choices^6^. By assigning values to options, the brain reduces complex, multidimensional decisions to a single dimension, facilitating efficient decision making^7^. The *orbitofrontal cortex* (OFC) is a key region supporting good-based decisions. Studies in non-human primates revealed three groups of neurons in OFC essential for economic choice: *offer value neurons*, encoding the value of individual options; *chosen value neurons*, representing the value of the selected option; and *chosen good neurons*, indicating the identity of the chosen good^6^. Because our model operates in good space without spatial contingencies, we refer to this last category as choice neurons. These neurons exhibit *menu invariance*, maintaining consistent encoding regardless of alternatives—a property supporting choice transitivity^8^. Electrical stimulation established a causal link between OFC neuronal activity and choice behavior supporting OFC’s integral role in the decision circuit^9,10^, though the division of labor between the OFC and adjacent prefrontal areas remains under investigation^11^.

The *circuit mechanisms* underlying *value computation* and *value comparison* are poorly understood^10,12^ and key questions remain open: How does a neural circuit, using additive synaptic inputs, implement the fundamentally multiplicative computation needed to calculate an expected value from probability and quantity? Where in the circuit are the learned relative values defining economic indifference points physically stored? And how are these value representations composed such that a single circuit flexibly performs multiple, diverse tasks? Moreover, existing studies often focus on single-neuron analyses and binary choice tasks, which don’t capture neural population dynamics or the complexity of real-world decisions involving multiple options and attributes^7^.

To gain mechanistic insights, computational modeling is essential^13–15^. However, previous models were often overly constrained, assuming strict segregation of neuronal roles and tuning properties not observed experimentally^16^, or too abstract, employing architectures like Gated Recurrent Units (GRUs) lacking clear biological counterparts^17^.

To bridge these gaps and provide concrete, testable hypotheses, we developed a biologically plausible computational model.

Our model is a continuous-time vanilla RNN (no GRUs) adhering to *Dale’s law* ^18,19^. Neurons have biologically realistic single-unit time constants. We trained the network using *Proximal Policy Optimization (PPO)* reinforcement learning^20,21^. This approach is more biologically plausible than supervised learning, mirroring animal training via reward feedback^21^. Furthermore, PPO’s training stability enabled us to use a simple vanilla RNN, avoiding less biologically plausible gated architectures.

Here, we show that networks trained on a diverse battery of economic choice tasks discover a two-stage computational organization. First, value computation occurs via feedforward learned input weights that physically store subjective preferences and approximate non-linear offer feature multiplication. This mechanism directly enables generalization to novel offers^22^. Second, value comparison is implemented within the recurrent circuit via a CRI motif mediating a robust winner-take-all decision^23,24^. Finally, we demonstrate that this architecture supports multitasking through the emergence of a compositional code combining a shared computational core with specialized neural modules^25,26^. Our model reproduces key neurophysiological findings from the primate OFC^6^ and provides a coherent, mechanistically grounded framework for the neural basis of economic choice.

## RESULTS

### Training neural networks for multiple choice tasks

To investigate neural mechanisms underlying economic decisions, we trained biologically plausible excitatory-inhibitory RNNs to perform a range of complex economic choice tasks (Figure 1A). These tasks required networks to compute and compare values in diverse contexts involving different goods, quantities, probabilities, and temporal structures.

**Figure 1.**
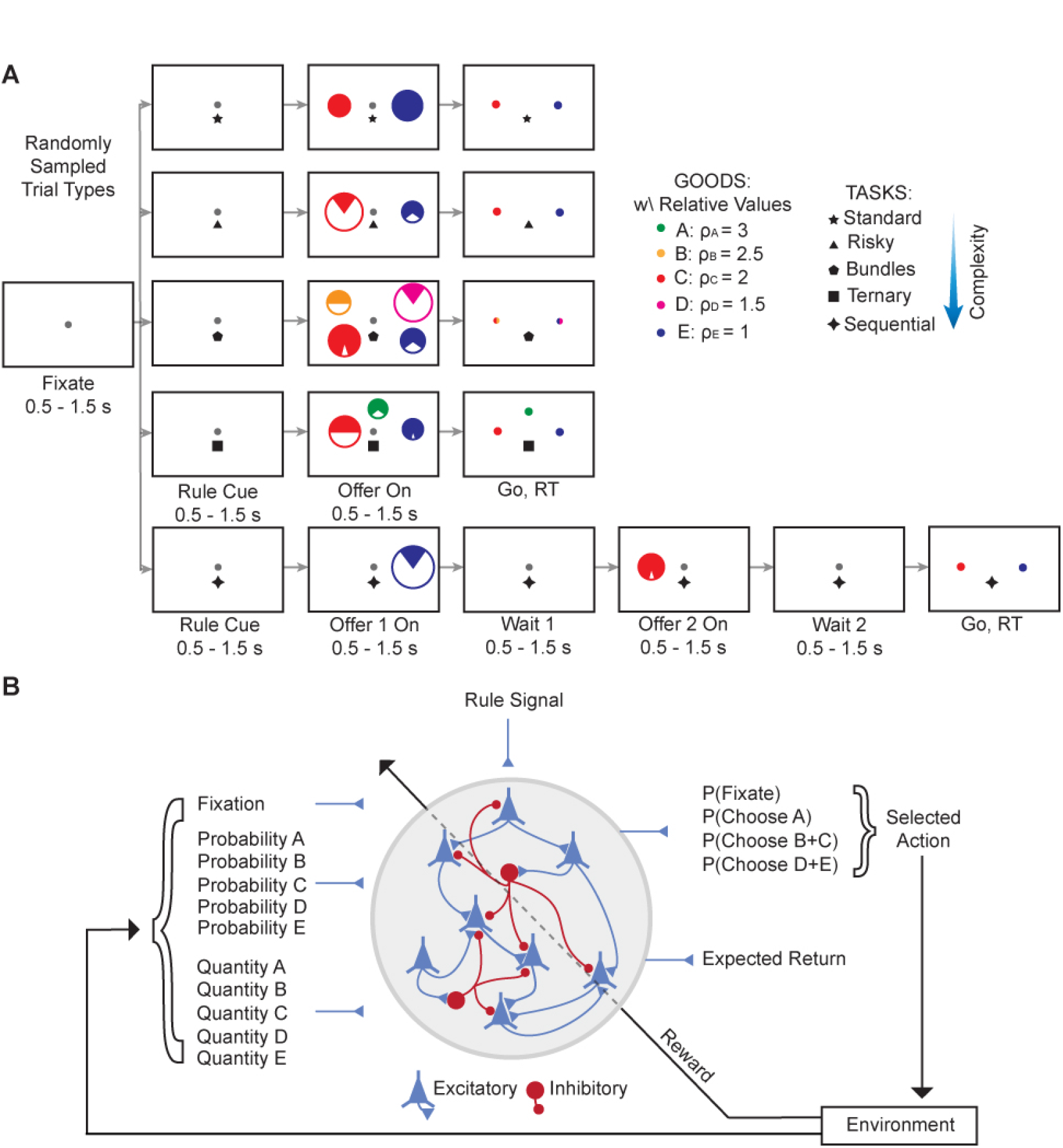
Task structures and network architecture. (A) Schematic of economic choice tasks. Trials begin with a fixation period, followed by a rule cue indicating the current task, offer presentation, and a response phase. Tasks include standard, risky, bundles, ternary, and sequential. For each offer, radius indicates quantity, filled area probability, and color corresponds to intrinsic value (A–E). (B) Biologically plausible RNN architecture with excitatory (E) and inhibitory (I) neurons adhering to Dale’s law. Inputs include fixation signals, quantities, probabilities, and task-specific rule cues. The network has two readouts: a policy output for action selection (the “actor”) and a value function predicting expected return (the “critic”). Training uses Proximal Policy Optimization in an agent-environment interaction loop.

Each task began with a fixation period, followed by a rule cue indicating the task, offers, and a response phase. Choice tasks included the *standard task*, where two goods were offered in varying quantities; the *risky task*, similar to the standard task but with probabilistic outcomes; the *bundles task*, offering bundles of two goods; the *ternary task*, involving choices among three goods; and the *sequential task*, where two goods were presented sequentially in random order, and choices relied on working memory^7^.

The networks consisted of continuous-time vanilla RNNs with 80% excitatory (E) and 20% inhibitory (I) neurons obeying Dale’s law, and long-range E projections (Figure 1B)^18,19,27^. Inputs included fixation signals, quantities and probabilities of offered goods, and task-specific rule cues. The networks produced two distinct outputs: a policy for action selection (the “actor”) and a value function predicting expected discounted future reward (the “critic”). The policy output represented probabilities over available actions at each time step, while the value function provided an ongoing estimate of expected return. This dual-readout architecture guided action selection and enabled value computation and evaluation.

We trained networks using PPO^20^, a reinforcement learning algorithm that simultaneously optimizes action selection and value estimation. This approach mirrors how animals are trained in laboratory experiments (i.e., through trial-and-error, with reward feedback^21^) and is thus more biologically plausible than supervised learning. Training involved an agent-environment interaction loop, where the network received inputs and selected actions leading to new stimuli and outcomes.

Networks achieved high performance across all tasks, satisfying two criteria: on a held-out test set of 25, 000 trials, they achieved *>* 99% trials completed without fixation breaks and selected the highest-value offer in *>* 90% of those trials (Figure S1).

We present sample trials for each choice task from a network trained simultaneously on all tasks (Figure S2). In each trial, the network selects the highest-value offer. The value function output predicts expected return shortly after offers are presented, demonstrating value computation. The policy outputs show correct action selection during the response phase and indicate fixation maintenance. Notably, the forthcoming choice can often be inferred from policy outputs during offer presentation, well before the response period. This suggests the network separates the economic decision (what to choose) from the motor command (when to respond).

In the sequential task (Figure S2E), the network received the first offer, maintained it in memory during the delay, and compared it with the second offer upon presentation. It then selected the higher-value offer, indicating effective integration of sequential information. This performance suggests the development of working memory capabilities essential for retaining and comparing offers over time.

These results demonstrate that our RNNs perform multiple complex choice tasks. Indeed, networks achieved high accuracy across all tasks, processing different types of information—quanti probabilities, and temporal sequences. Thus, networks developed the computational mechanisms for value-based decisions in diverse contexts.

### Networks learn a robust, value-based behavioral strategy

Choice data were analyzed via logistic regression^7^. In the risky task, trials involved choices between goods C and E, varying in quantity and probability. Plotting choices in quantity-probability space revealed no clear decision boundary (Figure 2A, left/center). However, fitting a logistic model to the choice data revealed a linear separation in the resulting *offer value space* (Figure 2A, right and 2B). Here, offer values are the product of quantity (*q*_*X*_), probability (*p*_*X*_) raised to a power 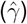, and intrinsic value 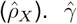 and 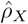 were key behavioral parameters estimated by the regression: 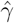 quantified risk attitude, and 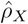 represented inferred relative value between goods (see STAR Methods).

**Figure 2.**
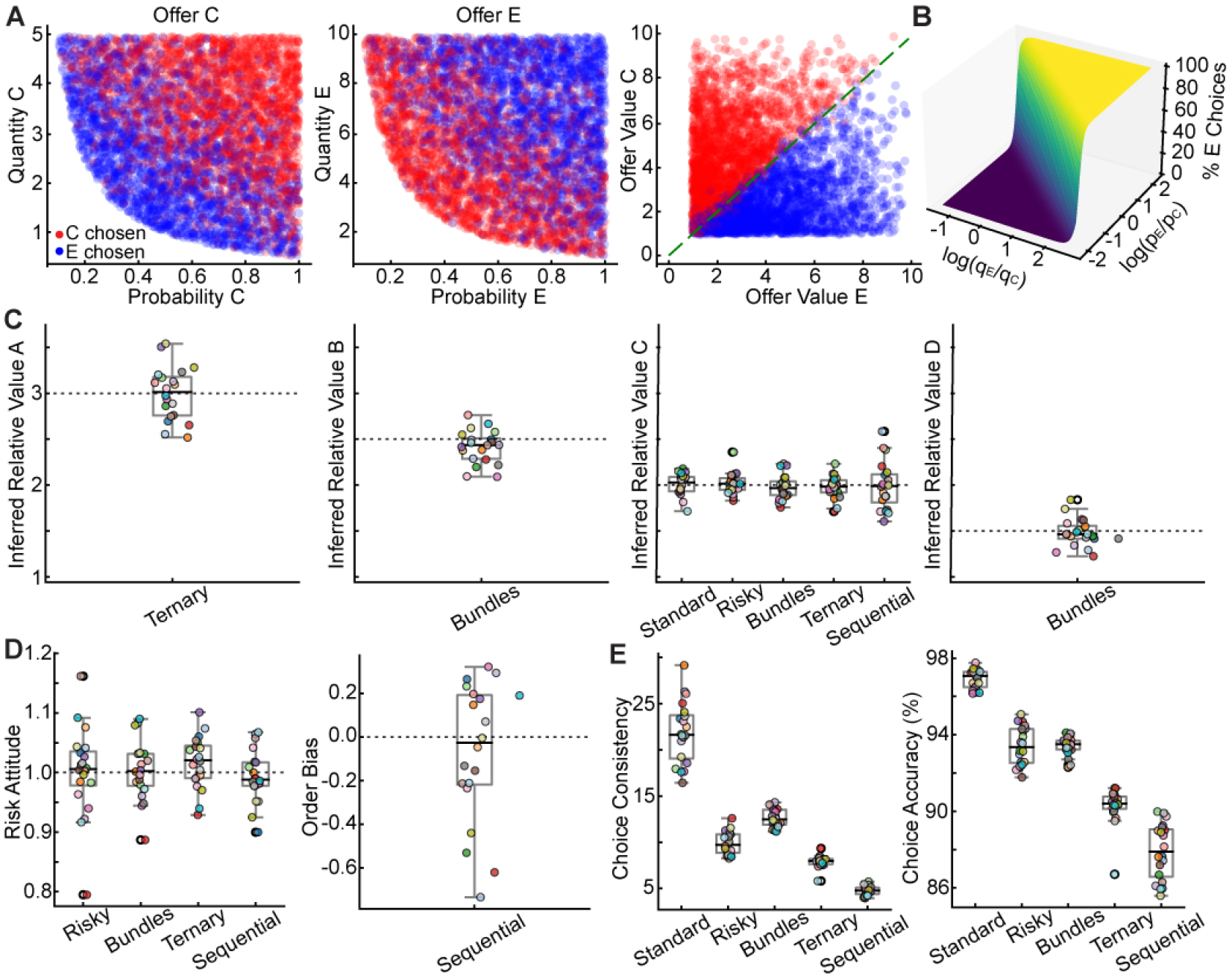
Logistic behavioral analysis of networks trained on all choice tasks. (A) Choice patterns in the *risky task* from a representative network. Left/Center: In the space of raw offer features (quantity and probability), network’s choices (red for C, blue for E) are not linearly separable. Right: When plotted in computed *offer value space* 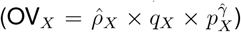 (parameters fit to behavior), same choices become clearly separable by the identity line, demonstrating a value-based strategy. (B) Psychometric function for the risky task (3D surface). The surface shows the percentage of choices for good E as a function of log-ratio of quantities (log(*q*_*E*_*/q*_*C*_)) and log-ratio of probabilities (log(*p*_*E*_*/p*_*C*_)). The steep sigmoidal shape demonstrates highly consistent, value-driven choices. (C) Inferred relative values 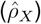 across all 20 networks (each point = one network) closely match ground-truth intrinsic values used in training (*ρ*_*X*_, dashed lines). The learned value ranking 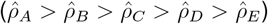 is robustly preserved. (D) Estimated behavioral biases are minimal on average. Left: The risk attitude parameter 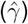 clusters around 1.0 (risk-neutrality), with minor variability. Right: Order bias (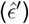 in the sequential task is centered at zero (no bias). (E) Estimated choice consistency 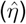 and accuracy vary systematically with task complexity. Per-formance is highest for the simple standard task and lowest for the more demanding sequential task, mirroring empirical findings.

Extending this analysis to the other tasks revealed a consistent strategy. In the standard task, networks compared computed values of two goods based on their quantities and intrinsic values, reliably selecting the higher-value good. Similarly, in the bundles task, networks computed the total value of each bundle by summing the values of individual goods to choose the higher total. In the ternary task, despite the complexity of comparing three options, networks reliably selected the highest computed value. In the sequential task, networks maintained the first offer value in working memory and compared it with the second offer value to make the optimal choice (Figure S3).

Logistic analysis allowed us to assess whether the network learned relative values from reward feedback alone. Treating relative values as free parameters estimated directly from choice behavior revealed a striking correspondence: *inferred relative values* 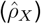 closely matched *ground-truth intrinsic values* (*ρ*_*X*_) used during training (Figure 2C). This mapping confirmed that networks learned the value structure of the environment to guide decisions.

We also examined *behavioral biases* including risk attitude and order bias. For probabilistic tasks (risky, bundles, ternary, sequential), we estimated the risk attitude parameter 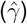 (Figure 2D). 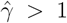 indicated risk aversion; 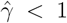 indicated risk seeking. While networks were risk-neutral on average 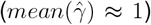, individual networks exhibited a range of mild risk preferences 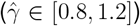. This variability likely arose from stochastic factors (e.g., random weight initializations, trial order of trials encountered during learning) as networks were trained to performance criteria without being overtrained. Notably, the risk attitude was consistent across tasks, suggesting a shared computational mechanism for processing risk—a key principle of compositionality that we revisit later. In the sequential task, we assessed the estimated order bias 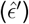—preference for first or second offer regardless of values. Again, while the average order bias across networks was negligible, individual networks exhibited slight preferences, reflecting stochastic fluctuations during learning.

Finally, we assessed the relationship between *task complexity and choice accuracy*. The logistic regression provided a measure of choice consistency 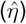 proportional to the slope of the psychometric function. We found that network performance scaled with task complexity: networks achieved higher choice consistency 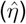 and accuracy on simpler tasks (e.g., standard task) and lower performance on more complex ones (e.g., sequential task; Figure 2E). This pattern aligns with empirical observations in non-human primates, suggesting that task difficulty impacts decision performance^7^.

To ensure robustness, we trained 20 independent networks on the full set of tasks. Despite different initializations, analysis revealed a remarkable consistency. While minor quantitative differences existed, logistic regression showed that all 20 networks converged on the same value-based strategy. All of them learned intrinsic relative values, exhibited comparable choice consistency, and developed modest idiosyncratic biases (Figures 2C-E). This convergence suggests that the computational mechanisms represent a robust solution. Whether different networks also converged on similar internal connectivity and dynamics is a central question addressed in the following sections.

### Single-neuron signatures of value computation and comparison

To investigate the neural code for economic choice, we first examined the activity of individual neurons. Across 20 independently trained networks, we consistently observed the same key functional cell types—offer value, chosen value, and choice neurons—indicating a convergent solution. The following analyses are therefore representative of a general coding strategy. We assessed whether individual neurons represented decision variables similar to those found in the primate OFC^6,28^. Focusing on binary choice tasks (standard, risky, sequential), we defined standard neurophysiological candidate variables: individual offer values (OVC, OVE), chosen value (CV), choice (CH), value sum (OVC + OVE), and value difference (OVC - OVE). We also defined two good-specific chosen values (CVC, CVE), which represented the value of a good (C or E) when that good was chosen and zero otherwise.

The correlation matrix (Figure 3A) revealed significant task structure. Chosen value correlated positively with the maximum offer values and with the value sum. Similarly, choice was strongly correlated with the value difference, as larger differences deterministically drive choice. The conjunctive variables CVC and CVE were correlated with their respective offer values and choices, reflecting their composite nature.

**Figure 3.**
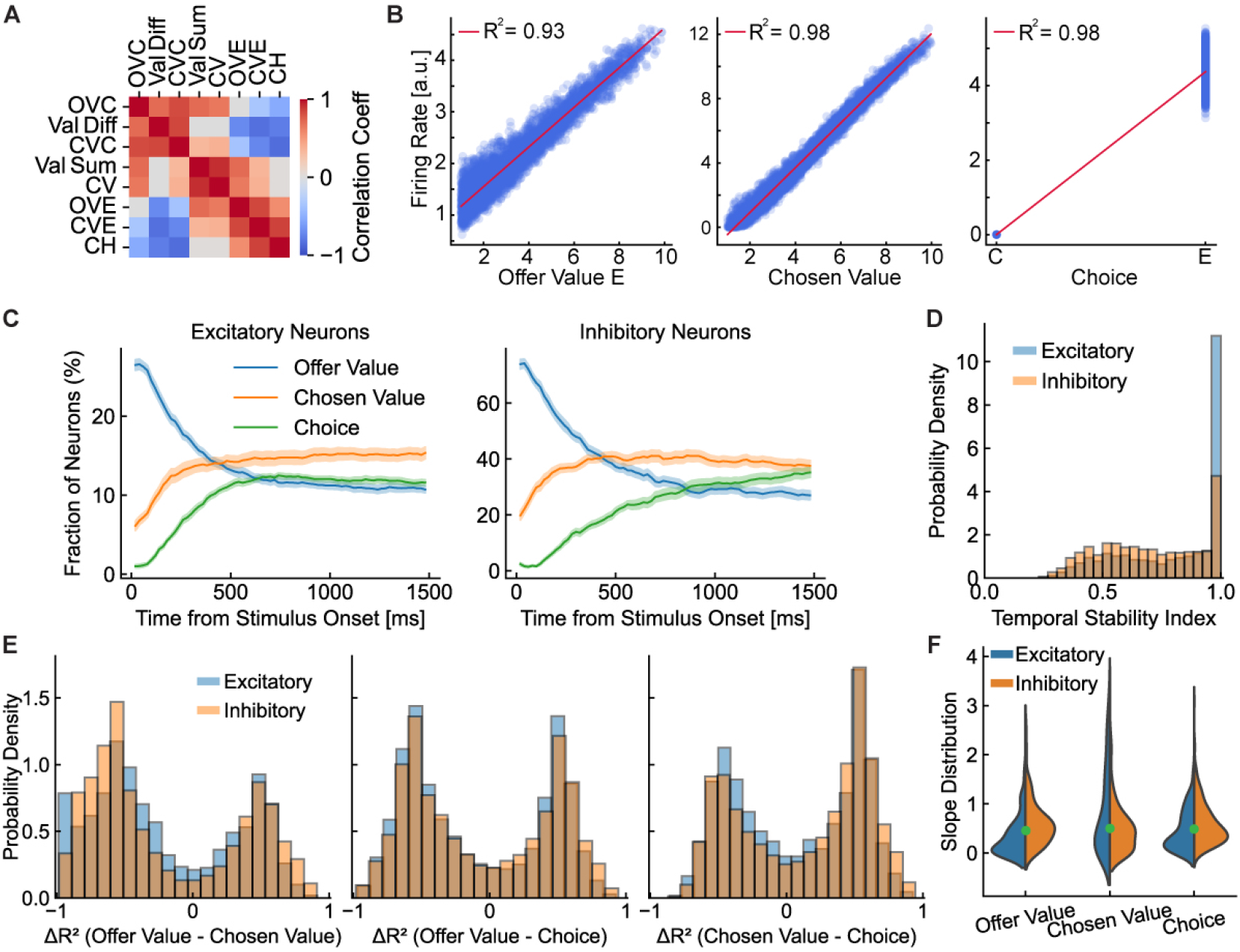
Single-neuron analysis in networks trained on all economic choice tasks. (A) Correlation matrix of behavioral variables for binary choice tasks. Reveals intrinsic task correlations, such as the positive correlation between chosen value (CV) and sum of offer values. (B) Example tuning curves from individual neurons in binary choice tasks. Activity predicts offer value of good E (OVE), chosen value (CV), and choice (CH). High coefficients of determination (*R*^2^ *>* 0.9) demonstrate highly specialized neurons. (C) Fraction of neurons selective for each decision variable over time. Using standard statistical criteria (*p <* 0.05, *R*^2^ ≥ 0.005), a substantial fraction of selective neurons appear in excitatory (left) and inhibitory (right) populations. Note that while the fraction of selective inhibitory neurons is high, the absolute number of selective excitatory neurons is larger due to the 80*/*20 E/I ratio. (D) Distributions of Temporal Stability Index (TSI). Most neurons show high TSI values, indicating stable encoding of a single decision variable throughout the trial. (E) Categorical encoding in a subpopulation of highly specialized neurons. Histograms show difference in *R*^2^ values between pairs of decision variables. Analysis includes only neuron-time points where at least one variable is explained with high efficacy (*R*^2^ ≥ 0.5). The resulting bimodal distributions suggest highly specialized neurons encode variables categorically. (F) Distribution of regression slopes for selective neurons. Both excitatory and inhibitory populations contain a mix of positively and negatively tuned cells for each key decision variable.

Analyzing tuning properties, we found that many neurons displayed significant linear relationships with specific decision variables. Figure 3B illustrates several examples. One neuron showed activity that increased linearly with the offer value of good E, another correlated with chosen value, and a third was selective for the binary choice. These neurons exhibited significant tuning (*R*^2^ *>* 0.9, *p <* 0.05; see STAR Methods).

To investigate the dynamics of neuronal selectivity, we calculated the fraction of neurons tuned to each decision variable at each time point (Figure 3C). Using standard statistical criteria for selectivity (see STAR Methods), we found a substantial fraction of neurons—peaking around 40–50% for key variables—were significantly tuned. This proportion was consistent with experimental observations^6^. The temporal evolution of this selectivity revealed a clear sequence that mirrored the decision process: during offer presentation, the fraction of neurons encoding offer value peaked first, followed by a rise in the fraction of neurons encoding the chosen value, and finally choice.

Interestingly, both excitatory and inhibitory neurons showed robust, dynamic selectivity. While the proportion of selective inhibitory neurons was higher (Figure 3C), given the network’s 80*/*20 E/I ratio, the total number of selective excitatory neurons was substantially larger. The high fraction of selective inhibitory cells aligned with their critical role in shaping circuit dynamics^29^.

To assess whether tuning was stable over the course of the trial, we computed a Temporal Stability Index (TSI) (Figure 3D; see STAR Methods). Most neurons showed high TSI, indicating consistent encoding of a single decision variable throughout the trial. However, some neurons showed lower TSI, suggesting a dynamic coding evolution from offer value to choice, unfolding during the decision process. This dynamic tuning aligned with experimental observations^28^.

We next investigated whether neurons encoded decision variables categorically or conjunc-tively. Paralleling analyses from primate electrophysiology^28^, we analyzed the distribution of *R*^2^ differences (Δ*R*^2^) between pairs of variables (Figure 3E; see STAR Methods). This revealed a dual coding scheme. While the general population of task-modulated neurons exhibited widespread mixed selectivity (see Supplementary Figure S4), a distinct subpopulation of highly specialized neurons developed categorical representations. When we isolated these strongly selective “expert” neurons by applying a stringent inclusion criterion, the resulting Δ*R*^2^ distributions became clearly bimodal (Figure 3E). Thus, “expert” neurons tended to encode one variable exclusively, allowing unambiguous representation of key decision variables.

Examining the sign of encoding, we found that both excitatory and inhibitory neurons displayed a mix of positive and negative correlations with their respective decision variables (Figure 3F). While a bias towards positive encoding was apparent, the emergence of a population of negatively tuned neurons was a critical feature. This bias was not related to valence of outcomes, as synaptic signs were fixed by Dale’s Law. Instead, it likely reflected the predominantly excitatory drive favoring positive tuning slopes. In contrast, negative tuning was a non-trivial emergent property of the learned recurrent circuitry. As shown later, this heterogeneity—especially negative encoding generated by targeted inhibition—was crucial for the competitive process. This diversity in tuning, including the bias towards positive slopes, was consistent with empirical findings^6,28,30,31^.

In conclusion, our single-neuron analyses reproduced key features of neuronal encoding observed in OFC. The sequential activation of neurons encoding offer value, chosen value, and choice reflected the computational stages of decision making. The active participation of inhibitory neurons and the presence of positive and negative tuning expanded our understanding of the mechanisms underlying value computation and comparison. These findings support economic decisions emerging from distributed computations within recurrent neural circuits, with neurons dynamically encoding relevant variables to guide behavior.

### Population-level signatures of value computation and comparison

To understand how neural populations collectively implemented economic choice, we analyzed their dynamics in state space.

We first quantified the complexity of neural representations across all tasks by measuring the embedding dimensionality of population activity using the participation ratio (PR). PR provided a continuous estimate of the number of dimensions a neural population used to represent information (see STAR Methods). The dimensionality of the manifold supporting neural activity was low for binary choice tasks (PR ≈ 2), and increased only for the ternary task (PR ≈ 3; Figure 4A). This pattern held for both excitatory and inhibitory populations. Low dimensionality suggested that networks efficiently organized their activity around the few variables necessary for the decision.

**Figure 4.**
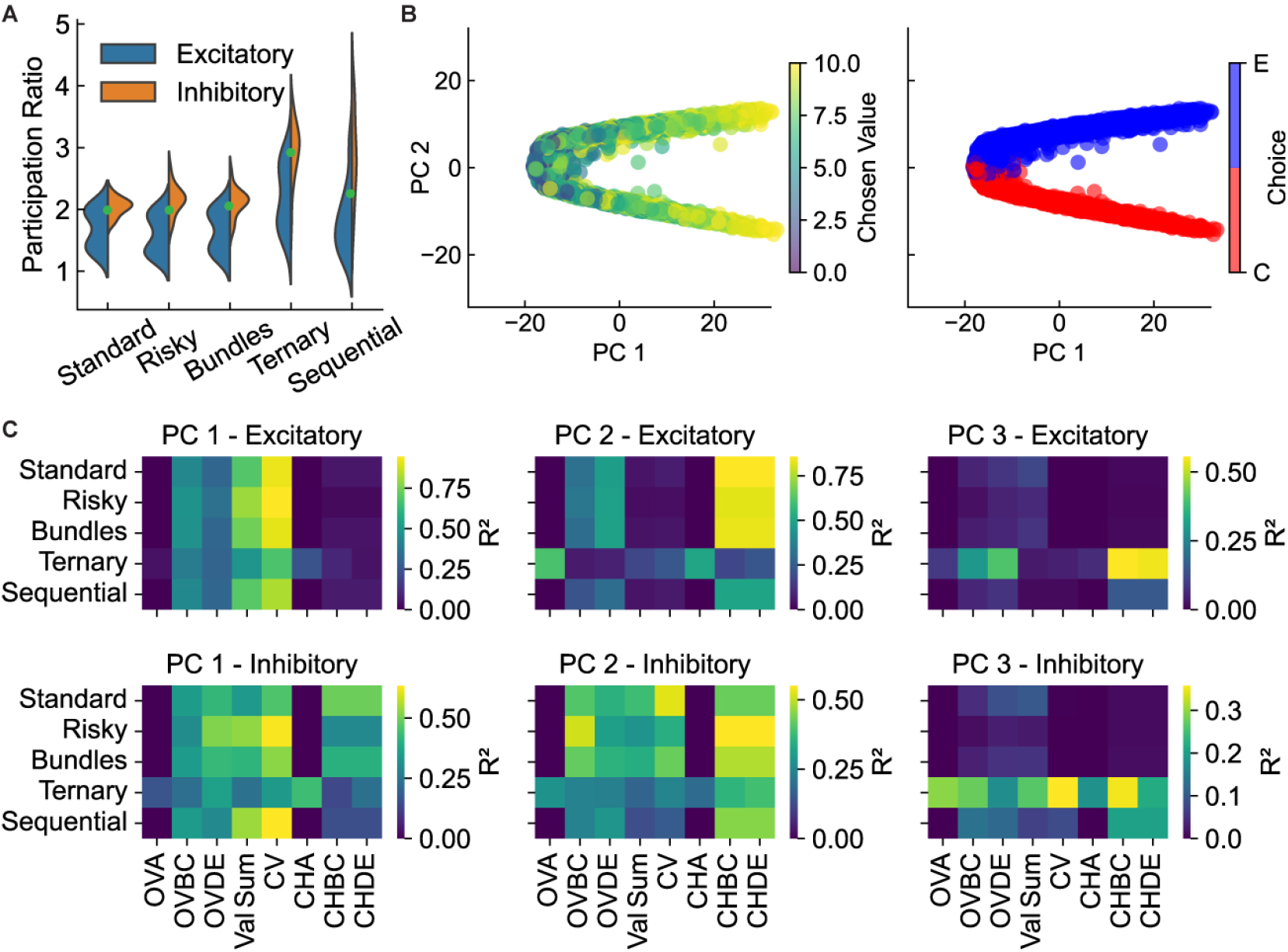
Low-dimensional population dynamics reveal a robust geometric solution for decision-making. (A) Neural dimensionality across tasks, quantified via participation ratio (PR). For both excitatory (blue) and inhibitory (orange) populations, neural activity was low-dimensional (PR ≈ 2), increasing to PR ≈ 3 only for the three-choice ternary task. (B) In the risky task, at the end of offer period, population states are geometrically organized by key decision variables. Each point represents the population state for a single trial projected onto the first two principal components (PC1, PC2). States are separated into distinct clusters based on choice (red vs. blue), with position within each cluster graded by chosen value. (C) Summary of population-level encoding across all tasks, quantified by *R*^2^ from regressing PC projections against key behavioral variables. X-axis: offer values for specific tasks (*OVA*: Offer Value of good A; *OVBC*: Offer Value of Bundle B+C; *OVDE* : Offer Value of Bundle D+E), a general value metric (*Val Sum*: sum of all offer values), and post-decision variables (*CV* : Chosen Value; *CHA, CHBC, CHDE* : categorical variables for Choice of good A, bundle B+C, or bundle D+E). Heatmaps reveal a consistent coding scheme: PC1 robustly encoded Chosen Value (CV), PC2 encoded Choice, and PC3 was selectively recruited for the ternary task.

At the population level, the decision manifested as a geometric separation of neural states. Using the risky task as an example, at the end of the offer period, the state of the network for each trial—projected onto the first two principal components—occupied a position in a highly structured cloud (Figure 4B). This cloud was organized by key decision variables: states were separated into distinct clusters based on the final choice, while the position within each cluster was graded by the chosen value. Supplementary Video S1 illustrates the full dynamic evolution. To systematically quantify this encoding, we regressed the principal component projections against all task variables (Figure 4C). Results revealed a consistent coding scheme: PC1 consistently encoded chosen value, while PC2 robustly encoded the choice itself. The role of the third principal component (PC3), however, was task-dependent. While largely un-encoded in binary tasks, PC3 became critical for representing the third option in the ternary task (Fig. 4C, third column). This demonstrated an efficient coding strategy: a core, two-dimensional geometric solution for value and choice was conserved across all contexts, and the network flexibly recruited additional dimensions only as required by the specific demands of the task.

Importantly, this emergent geometric solution was not an idiosyncratic feature of a single network: The same separation of population states into choice-specific clusters organized by chosen value was a convergent strategy in all 20 networks (see Supplementary Figure S5).

### Dissecting the circuit mechanisms of value computation

Targeted removal of recurrent connections (*W* ^rec^ = 0) revealed a functional dissociation: while lesioned networks could not choose, their feedforward architecture preserved value-related information (Supplementary Fig. S6, S7). This established value computation as primarily a feedforward process, while comparison is a distinct, competitive process reliant on the recurrent circuit. We next dissected how the feedforward pathway solves two problems: integrating offer features and storing subjective preferences.

First, how does a circuit endowed only with additive synapses approximate the non-linear *multiplication* of quantity and probability?

Activity in the isolated feedforward pathway was explained significantly better by the multiplicative product of quantity and probability than by their sum (Fig. 5B). In other words, the network approximates a non-linear, multiplicative computation.

**Figure 5.**
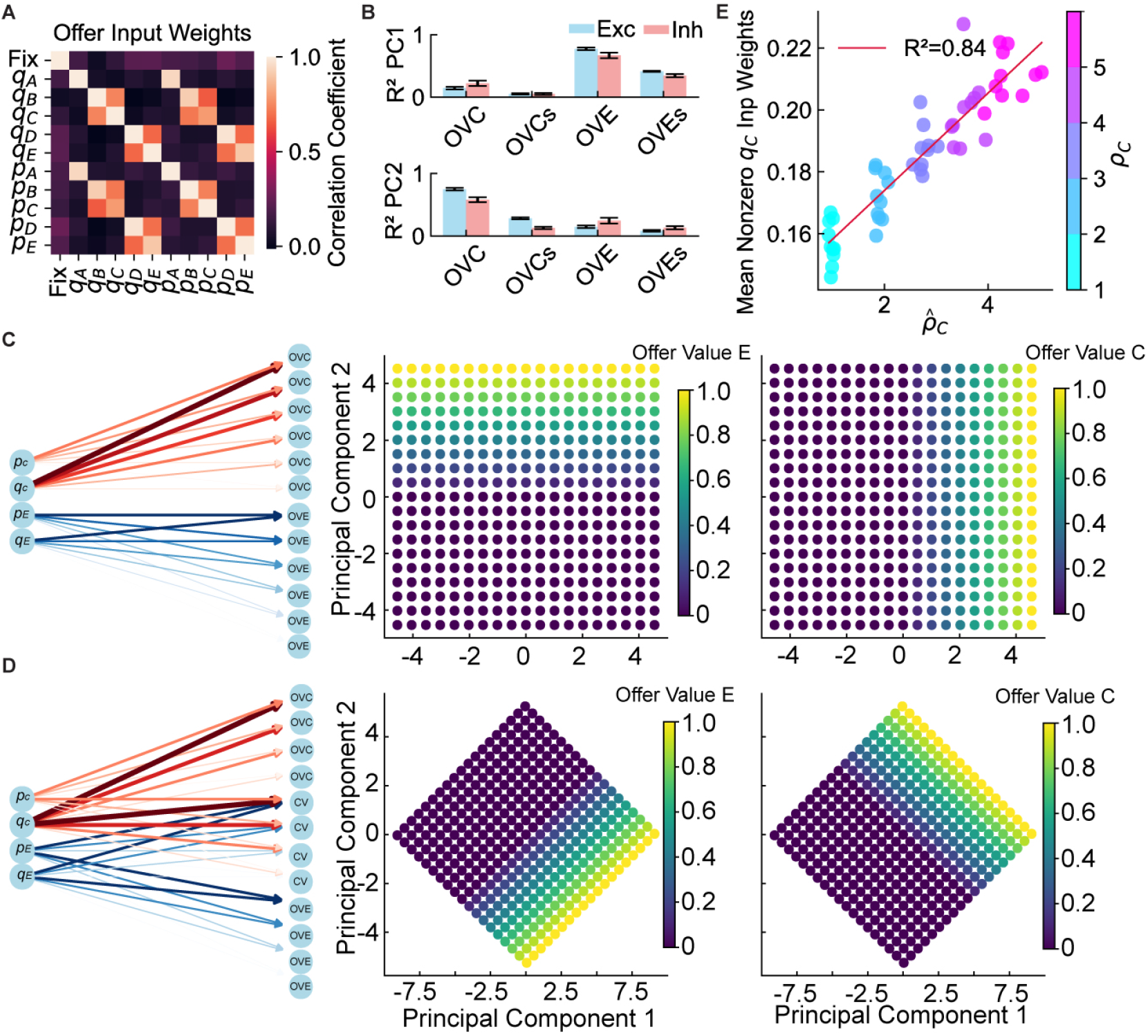
Dissecting the circuit mechanisms of value computation. (A) Correlation matrix of input weight vectors for offer features. Strong positive correlations (bright colors) appear between weights for features of the same offer (e.g., quantity C, *q*_*C*_, and probability C, *p*_*C*_), with weaker correlations across offers. (B) Evidence for multiplicative computation in the isolated feedforward pathway. In networks where recurrent connections were lesioned, the first two principal components (PCs) of population activity were better explained by the multiplicative product of quantity and probability (OVC, OVE) than by their additive sum (OVCs, OVEs). Bars show mean coefficient of determination (*R*^2^) across 20 networks; error bars denote SEM. (C) A simplified feedforward toy model with segregated input populations failed to reproduce the geometric rotation observed in trained lesioned networks (Supplementary Fig. S7B). PCA of hidden units’ activity revealed principal axes corresponding to individual offer values (Offer Value C on PC1, Offer Value E on PC2). (D) Extending the toy model with a population of mixed-selective neurons causally reproduced the rotation of principal component axes to align with value sum and difference. This validated the hypothesis that mixed-selective wiring (Supplementary Fig. S9) explains this geometric fea-ture. (E) Learned relative values are physically encoded in input weights. We trained 50 independent networks while systematically varying the ground-truth intrinsic value of good C (*ρ*_*C*_) across five groups (from 1 to 5). The plot reveals a strong, significant linear correlation (*R*^2^ = 0.84) between the average learned input weight for that good’s quantity (y-axis) and the behaviorally inferred relative value 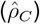, estimated from each network’s choices (x-axis). The color of each point indicates the ground-truth intrinsic value (*ρ*_*C*_) used to train that network. Thus, networks trained with a higher intrinsic value (brighter magenta points) not only developed stronger corresponding input weights but also learned to choose in a way that accurately reflected this value, as the inferred behavioral values 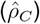 clustered correctly around their ground-truth counterparts.

The network achieved this by developing a function approximation over the domain of its experience, a hallmark of neural networks with non-linear activation functions, rather than by learning abstract, symbolic rules. Importantly, while this dissection analysis provided strong evidence for the mechanism’s capability, a stronger test for a learned computational principle required demonstrating its use by the intact network to solve novel challenges (see below).

Second, where are learned subjective *relative values* (*ρ*_*X*_) physically stored? We posited that these behavioral parameters are encoded in synaptic strengths of input projections. To test this hypothesis, we trained 50 independent networks on the risky task, varying one good’s intrinsic value across networks. A strong, significant linear correlation emerged between behaviorally inferred relative values and average learned input weights for that good’s quantity (Fig. 5E; *R*^2^ = 0.84). This confirmed that learned economic preferences were physically embedded in synaptic efficacies of long-range inputs. Such design allows generalization to novel goods without retraining recurrent decision circuitry, and thus is computationally efficient.

Finally, why does population geometry in the feedforward pathway rotate to represent the *value sum and value difference*, rather than individual offer values (Supplementary Fig. S7B)? We hypothesized that this geometry is optimized for the downstream decision process. A representation based on sum and difference is computationally advantageous for downstream readout, as the chosen value (the primary variable for critic and actor) is highly correlated with the value sum under the uniform sampling distribution. To identify the structural basis for this organization, we examined the correlation structure of the input weights, which revealed a mix of specialized and integrated pathways for the different offer features (Fig. 5A). To directly quantify this wiring scheme, we performed a Selectivity Index (SI) analysis on the input weights. This revealed a dual architecture (Supplementary Fig. S9): in addition to two populations of highly specialized neurons that exclusively processed a single offer’s value (corresponding to the sharp peaks at SI ∈ {− 1, +1 }), we found a large, unimodal population of mixed-selective neurons receiving balanced inputs from both offers (the broad distribution centered at SI = 0). To verify this, we constructed a simplified feedforward “toy model”. A model with only segregated input populations failed to reproduce the rotated geometry, instead representing offer values on orthogonal axes (Fig. 5C). However, after incorporating a population of mixed-selective neurons, the model successfully replicated the value sum/difference representation (Fig. 5D). This demonstrates that the networks learned a specific, non-trivial input wiring scheme explicitly structured to facilitate downstream decisions.

Thus, the feedforward pathway performed sophisticated value computations, integrating external features with internal (learned) preferences via a specific connectivity structure optimized for the downstream decision.

### Dissecting the circuit mechanisms of value comparison

Next, we investigated how the recurrent circuit *compares* values to generate a decision. To distinguish the computational outcome—a winner-take-all (WTA) competition—from the circuit mechanism implementing it, we analyzed learned connectivity and dynamics. We found that the WTA outcome relied on a specific E/I motif we termed Competitive Recurrent Inhibition (CRI). We causally validated this motif through lesion analyses testing the roles of its key constituent parts.

We first analyzed policy output units during the risky task (Fig. 6A). These units, representing available actions distinct from recurrent neurons, displayed a competition well before the go cue. During stimulus presentation, the “Fixation” output was dominant, gating the motor response (Fig. 6A, first panel). Meanwhile, a subtle competition emerged between choice options. When choosing good C (red traces), its corresponding output showed higher activity progressively diverging from the output for good E (Fig. 6A, third vs. fourth panels), and vice versa when choosing good E (blue traces). Thus, the network internally committed to a choice long before response execution. This early commitment was corroborated by reaction times (RTs) being independent of decision difficulty (Supplementary Fig. S10A). In contrast, the critic’s “Expected Return” output robustly encoded chosen value, independent of choice identity (Fig. 6A, fifth panel). This WTA dynamic at the output level was confirmed by regression analysis, showing that the choice output’s activity was predictive of the corresponding offer’s value only in trials where it was chosen (Fig. 6B).

**Figure 6.**
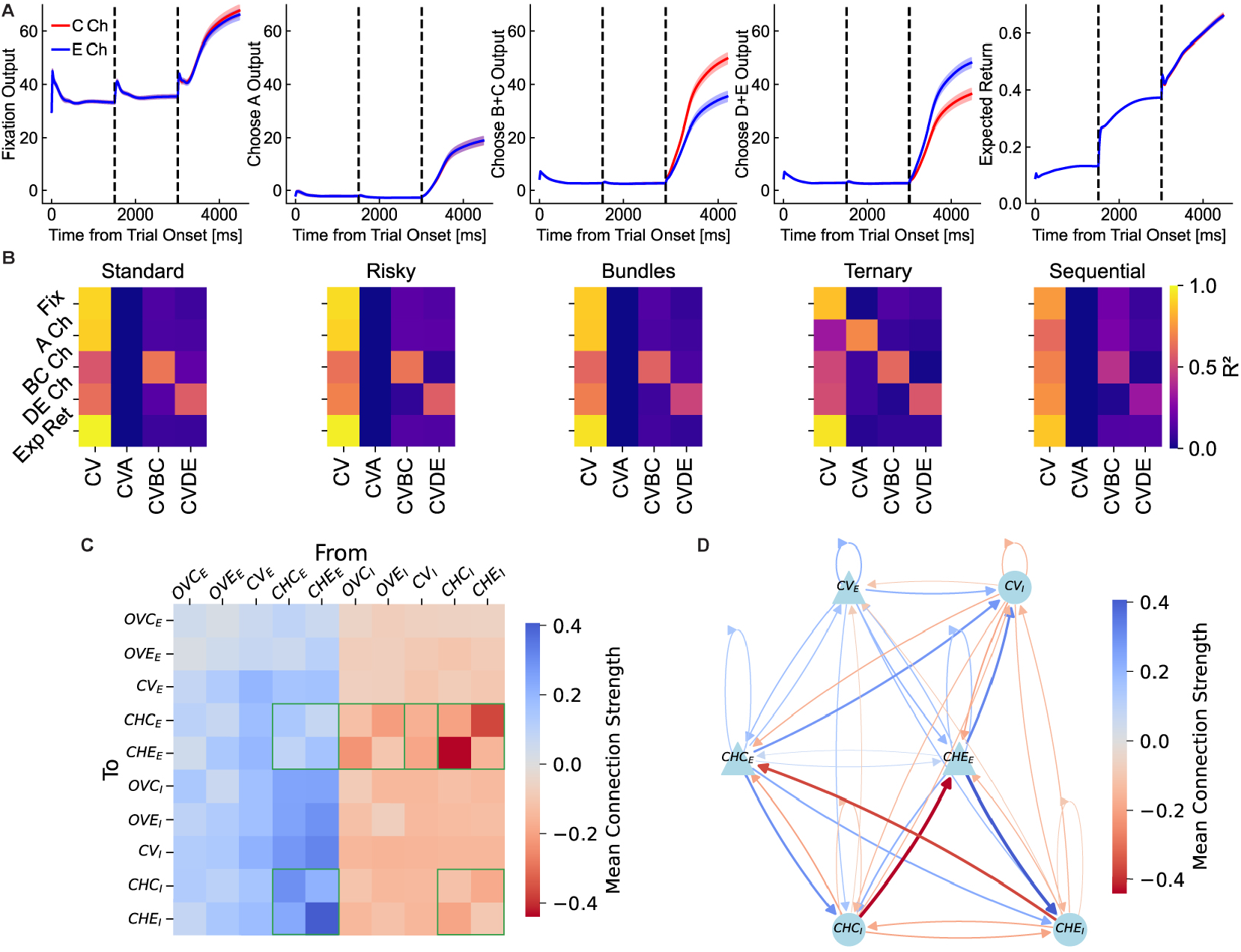
Dissecting the neural circuit mechanisms of value comparison. (A) Average activity of policy (actor) and value (critic) output nodes during the risky task, separated by final choice (red for chosen C, blue for chosen E). From left to right: fixation output, an irrelevant choice output, choice C output, choice E output, and critic’s expected return output. Divergence between red and blue traces for choice outputs indicates early commitment. (B) Heatmap of regression *R*^2^ values, confirming the WTA dynamic. Each panel corresponds to a different task. Within each heatmap, a row represents one of the network’s output nodes (e.g., ‘BC Ch’ is the policy output for choosing good C or B+C). Columns represent key decision variables, including general variables like Chosen Value (‘CV’) and conjunctive, choice-specific variables like ‘CVBC’ (the value of good C or B+C on trials where C or B+C was chosen, and zero otherwise). Each cell shows *R*^2^ from a simple linear regression of a single output node’s activity against a single variable across all trials. Analysis reveals a clear WTA pattern: the activity of a choice-specific output node was strongly predicted by its corresponding conjunctive value variable, but not by the variable for the competing choice. For instance, in the ‘Risky’ task, the ‘BC Ch’ output was well-predicted by ‘CVBC’ (bright square), but poorly predicted by ‘CVDE’ (dark square). This demonstrates selective encoding of the chosen option’s value at the actor level. (C) Reduced connectivity matrix, averaged across 20 networks. This was derived by averaging synaptic weights between functionally defined neural pools (see Supp. Fig. S11). The matrix revealed the Competitive Recurrent Inhibition (CRI) motif, highlighted by strong inhibitory connections (dark blue squares, outlined in green) from choice-selective inhibitory neurons to competing choice-selective excitatory neurons. (D) A simplified circuit diagram visualizing the CRI motif. Choice-selective excitatory neurons (e.g., CHE_*E*_) drove same-choice inhibitory neurons (e.g., CHE_*I*_), which in turn suppressed excitatory neurons corresponding to the competing choice (e.g., CHC_*E*_), implementing WTA competition. Acronyms: OVC/E (Offer Value C/E), CV (Chosen Value), CHC/E (Choice C/E). Subscripts _*E*_ and _*I*_ denote excitatory and inhibitory populations, respectively.

The low-rank structure the full 256 *×* 256 recurrent weight matrix (Supplementary Fig. S10B) suggested that the network’s dynamics were governed by few connectivity motifs.^32^ Thus to identify the circuit motif responsible for WTA competition, we computed a reduced, interpretable circuit diagram. We developed a *many-to-one functional abstraction* pipeline (Supplementary Figure S11). This process involved fixing a task (e.g. risky), classifying each neuron by cell type (E/I) and functional selectivity, grouping neurons in functional pools, and averaging connection strengths. Averages across all 20 networks revealed the CRI motif (Figure 6CD). In essence, choice-selective E neurons drove same-choice I neurons, which preferentially suppressed E neurons selective for the competing choice. For instance, ‘Choice C’ E neurons strongly recruited I neurons, which powerfully suppressed ‘Choice E’ E neurons.

This targeted, cross-choice inhibition mechanism shed light on two key findings. First, it directly implemented WTA competition. Second, it explained the emergence of heterogeneous tuning. While feedforward inputs were excitatory, targeted recurrent inhibition from the CRI motif could powerfully subtract from a neuron’s input drive, effectively inverting response profiles to create negatively encoding cells that sharpened contrast between choice representations. We hypothesized that these negatively tuned neurons were computationally critical. Indeed, silencing all neurons with significant negative encoding of offer value and chosen value substantially impaired decision-making across all tasks, significantly reducing choice accuracy (Supplementary Fig. S12). Thus, negatively tuned neurons – a direct consequence of the CRI architecture – were essential for robust value comparison.

Finally, to causally validate the CRI motif necessity, we performed a “functional ablation” analysis. We silenced entire functional pools of neurons and evaluated performance in the risky task. Our results revealed a distributed role for choice-selective populations (Supplementary Fig. S13). While silencing choice-selective inhibitory neurons ‘*CHE*_*I*_’ had the largest impact on trial completion, choice accuracy relied on a much broader set of neurons. Critically, silencing choice-selective inhibitory neurons (‘*CHC*_*I*_’, ‘*CHE*_*I*_’) and chosen-value excitatory neurons (‘*CV*_*E*_’) had effects on accuracy as large, or even larger, than silencing their excitatory choice counterparts. In other words, both excitatory and inhibitory components of choice-selective populations formed the functional backbone of the decision circuit.

This CRI motif was found across all choice tasks (Supplementary Fig. S10C), suggesting a general strategy for competitive selection. In summary, our findings indicate that the recurrent network performs value comparison through the CRI mechanism—structured recurrent connectivity implementing WTA dynamics and generating heterogeneous neuronal tuning to increase contrast between competing offers.

### Compositionality in economic decisions

A key feature of intelligent behavior is the ability to flexibly solve multiple tasks by composing a repertoire of learned skills. To investigate if single neural circuits could develop such compositional representations for economic choice, we analyzed 20 networks trained on all five tasks simultaneously. We sought to understand how the network balanced shared neural resources against task-specific computations.

First, we examined neural subspaces activated during the rule cue period. Low-dimensional subspaces for the standard, risky, bundles, and ternary tasks largely overlapped (small principal angles) (Fig. 7B). This shared geometry, consistent with the highly correlated rule input weights (Fig. 7A), meant that a dimensionality-reduction basis found for one task would generalize well to the others. This clarifies the explicit task cue’s role: for these four tasks, it becomes largely redundant after training because the network developed a shared internal context. However, for the sequential task, the cue remained critical to signal its unique temporal structure and engage working memory circuits. Analysis revealed two key specializations: the sequential task occupied a distinct, nearly orthogonal subspace, reflecting its unique working memory demands, while the ternary task, though overlapping, required an additional dimension to represent the third option (Fig. 4A). Together, these results show that the network developed a core set of shared dimensions for common computations while flexibly allocating specialized subspaces for unique task demands.

**Figure 7.**
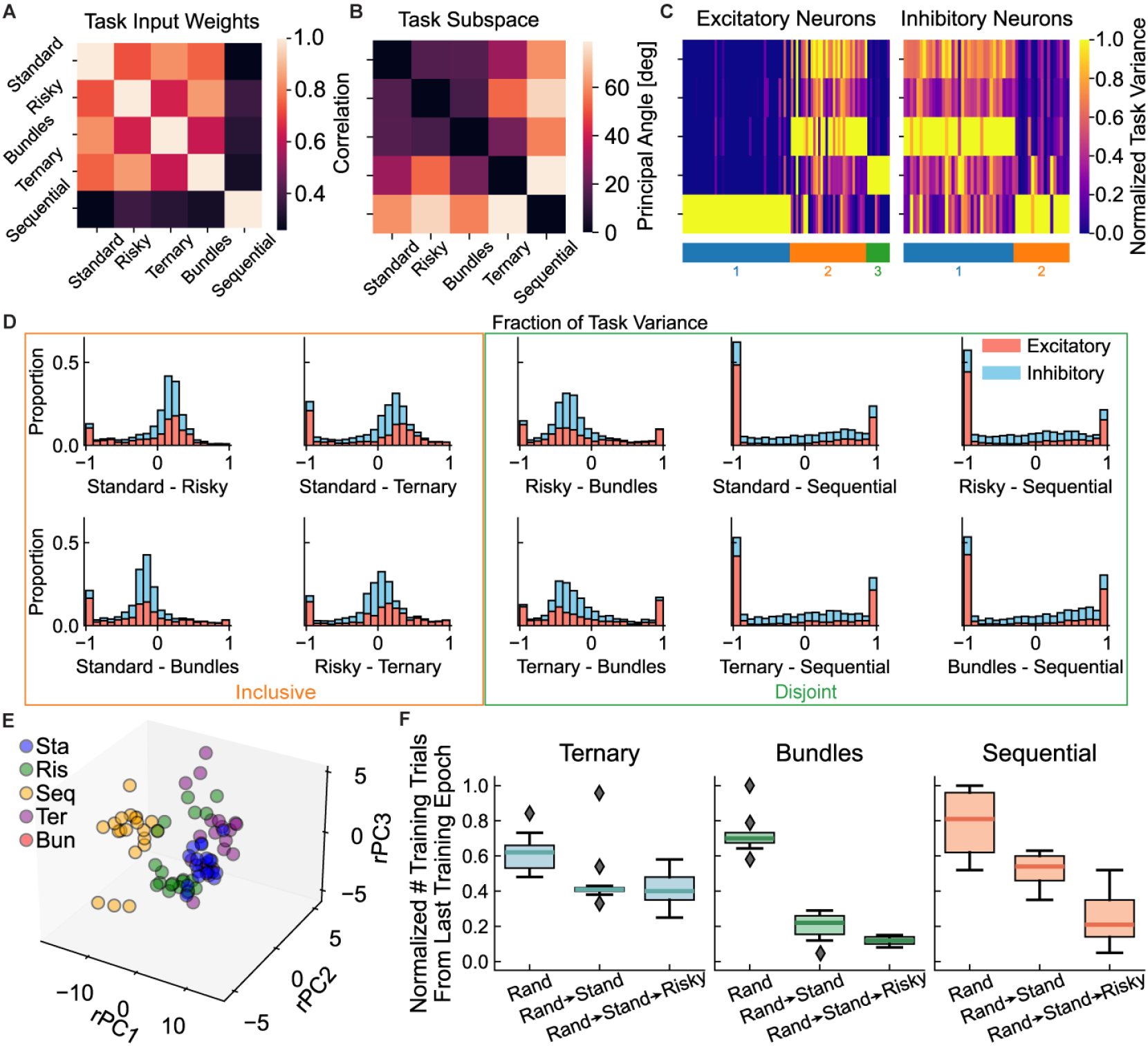
Compositionality and curriculum learning in networks trained on all tasks. (A) Correlation matrix of rule cue input weights. High correlation (light colors) among standard, risky, bundles, and ternary tasks indicates a shared input structure, distinct from the sequential task. (B) Subspace analysis of population activity during the rule cue period. Heatmap shows principal angles between low-dimensional subspaces for each pair of tasks. Small angles (dark colors) between subspaces for standard, risky, bundles, and ternary tasks indicate a common representational geometry. The sequential task occupied a distinct, nearly orthogonal subspace (bright colors). (C) Clustering of neurons by task variance reveals shared and specialized modules. Neurons (columns), separated by excitatory and inhibitory populations, were grouped based on their normalized activity variance across tasks (rows). Analysis identified distinct clusters, including a shared module active across all tasks, a specialized module with high variance primarily in the ternary task, and another specialized module almost exclusively dedicated to the sequential task. (D) Histograms of fractional task variance differences between task pairs, calculated as (*TV*_1_ − *TV*_2_)*/*(*TV*_1_ + *TV*_2_) for each neuron. The shapes of these distributions reveal organizational relationships between neural populations recruited for different tasks. Following Yang et al.^25^, we characterized these shapes, which revealed patterns resembling archetypal *inclusive* relationships (e.g., Standard-Bundles) and *disjoint* relationships (e.g., Ternary-Sequential). (E) Visualization of task representations in a common, rotationally-aligned PCA space aggregating data from all 20 networks. Each point represents mean population activity for one task in one network. The four simultaneous-offer tasks (Standard, Risky, Ternary, Bundles) formed a tight cluster, while the Sequential task formed a separate cluster, highlighting its distinct neural dynamics. (F) Curriculum learning accelerated skill acquisition. Box plots show the normalized number of training trials required to learn only the *final* complex target task (Ternary, Bundles, or Sequential) in each curriculum. For each target, learning speed was compared across three conditions (10 networks each): from random initialization (Rand), after pre-training on the Standard task (Rand→Stand), or after pre-training on Standard and then Risky tasks (Rand → Stand → Risky). Pre-training provided a significant, graded acceleration in learning the final task, demonstrating schema reuse.

While sharing a common choice structure, these tasks imposed unique computational demands testing true compositionality. For example, the *ternary task* required scaling WTA competition from two to three options, while the *sequential task* required to hold a value in working memory over time. To test whether networks developed distinct modules to meet these demands, we clustered neurons based on their firing rate variance across tasks, analyzing excitatory and inhibitory populations separately (Fig. 7C). This analysis revealed a similar functional organization in both cell populations, identifying principal clusters: a population shared across all tasks, a cluster specialized for the ternary task, and a cluster exclusively dedicated to the sequential task. To causally test the function of these identified modules, we performed targeted lesion analyses. We silenced each cluster individually and re-evaluated performance across all tasks. The results revealed a clear functional dissociation (Fig. S14). Lesioning the sequential-task cluster specifically impaired the sequential task while leaving other tasks largely intact. Similarly, lesioning the ternary-task cluster selectively impaired the ternary task. Conversely, lesioning the shared cluster caused profound deficits in all five tasks. This provided direct causal evidence that the network developed a compositional architecture of both shared and functionally specialized modules to meet unique task demands.

We further quantified task relationships by examining distributions of task variance differences for each neuron across task pairs (Fig. 7D; see STAR Methods).

Following established methods^25^, we classified these distributions to characterize the organizational relationship between neural populations recruited for different tasks. Analysis revealed a spectrum of relationships, including patterns resembling archetypal *inclusive* relationships (where one task’s neural representation is a subset of another’s, characterized by a large peak at zero and smaller peak sat *±*1) and *disjoint* relationships (where tasks recruit largely separate sets of neurons, identified by bimodal distributions with peaks at *±*1). This supported a flexible network architecture balancing common processing with task-specific adaptations.

To visualize whether all 20 networks converged on a similar compositional architecture, we compared their population-level representations. Notably, direct comparison of principal components is not meaningful, as each independently trained network develops its own arbitrary principal component axes—akin to its own private coordinate system. To overcome this issue, we used Procrustes analysis (STAR Methods) to rotationally align each network’s state space onto a common reference frame, effectively translating all private coordinate systems into a single, shared language. Projecting the mean population activity for each task into this aligned space (Fig. 7E) revealed a robust, convergent solution: across all networks, the representations for the four simultaneous-offer tasks consistently clustered together, while the sequential task formed a separate, distinct cluster, graphically underscoring the unique neural dynamics required for working memory.

A critical question was whether this compositional architecture was a unique product of the multitasking paradigm, or if single-task training would suffice. To address this, we performed a systematic “zero-shot” generalization analysis. We trained five separate groups of networks (10 per group) on each of the five tasks individually and tested performance on all other tasks without further training (Fig. S15). The results revealed strongly asymmetric generalization causally demonstrating the compositional nature of the learned skills. For example, networks trained on the complex ternary task could immediately perform the simpler standard and risky tasks with high accuracy. Conversely, networks trained only on the standard task failed catastrophically on the sequential task, lacking a learned working memory component. This analysis confirmed that while multitasking is not required to solve any single task, it is a training condition enabling a single circuit to develop a flexible repertoire of reusable computational modules.

Finally, we tested whether the network could leverage prior experience to accelerate the acquisition of more complex skills. We designed a curriculum learning protocol for three complex target tasks: Ternary, Bundles, and Sequential (Fig. 7F). For each target, we compared learning speed under three conditions: (1) training from random initialization (Rand), (2) pre-training on the simpler Standard task, and (3) pre-training on the Standard then Risky task before learning the target (*N* = 10 each). This design allowed us to isolate how learning foundational components (e.g., temporal structure from the Standard task, then probabilistic integration from the Risky task) facilitated acquiring a final, complex skill. The additional experience from pretraining was not a confound but the experimental variable of interest—the proposed mechanism for schema formation. The critical question was whether prior experience provided a useful computational scaffold accelerating future learning. The results showed a clear affirmative (Fig. 7F); for each target task, there was a graded and significant acceleration in learning, with the number of trials required to learn the *final* task decreasing at each stage of pre-training. This demonstrated the formation of a cognitive “schema”—a reusable set of computational components transferred to the new context to facilitate subsequent learning.

In summary, networks trained for multitask proficiency developed a compositional architecture combining a shared circuit core with functionally specialized, task-specific modules. The formation of these reusable schemas supported efficient learning and flexible decision making across diverse contexts.

### Multiplicative value computation enables generalization to novel offers

A hallmark of intelligent behavior is the ability to generalize learned knowledge to new situations. To validate our hypothesis that networks solve economic choice via feedforward functional approximation of multiplication, we challenged networks trained on a limited subset of experiences to infer the general valuation rule, applying it to entirely novel offers^22^.

We trained 10 networks exclusively on the risky task using a constrained, orthogonal offer set. In this regime, quantity and probability never varied simultaneously, forcing independent learning of their contributions (Fig. 8A; see STAR Methods). After training, networks performed with high accuracy on this familiar, constrained set. We then assessed generalization using a full, unconstrained set of offers, including novel quantity-probability combinations (Fig. 8B).

**Figure 8.**
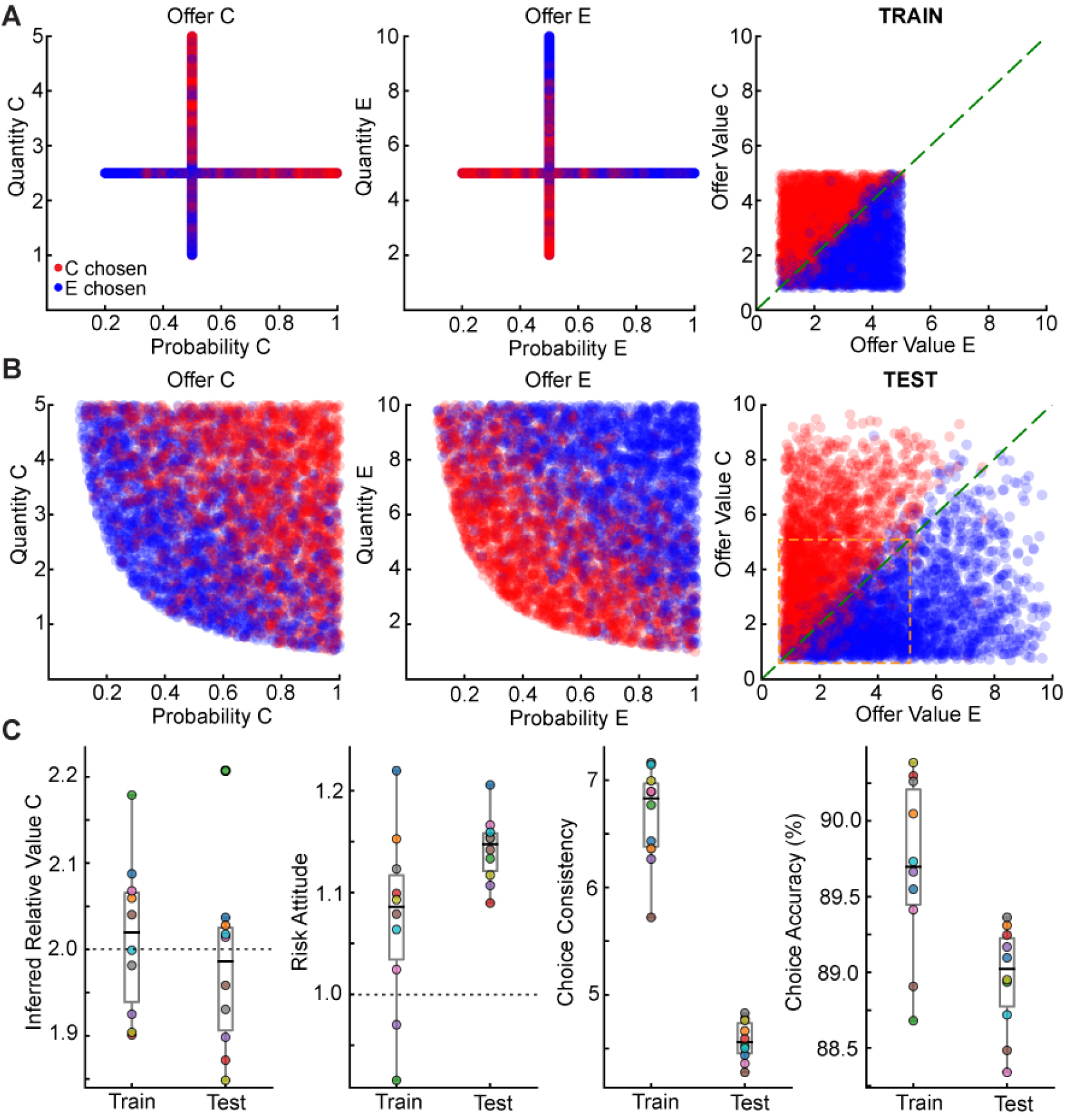
Multiplicative value computation enables robust generalization to novel offers in networks trained on a constrained risky task. (A) Training used a constrained set of offers where quantity and probability were not varied simultaneously, forcing independent learning of their contributions. Left/Center: The offer space for goods C and E during training forms a cross-like structure. Each point is a trial, colored by choice (red for C, blue for E). This visualization shows that the network was never exposed to trials where both quantity and probability varied from their central values. Right: Despite constrained inputs, choices plotted in computed offer value space are correctly and linearly separated by the identity line (dashed green), indicating successful learning. (B) Generalization tested via a full, unconstrained set of offers containing novel combinations of quantity and probability. Left/Center: The test offer space covers the full range of quantity-probability combinations, most never seen during training. Right: The network correctly evaluates these novel offers, as its choices remain well-separated by the identity line. This demonstrates successful generalization of the learned value function to unseen regions of the offer space. (C) Key behavioral parameters are stable between training and testing, confirming application of the same valuation strategy to novel stimuli. Each colored dot represents one of ten trained networks. Box plots show median and interquartile range. *Inferred Relative Value C* and *Risk Attitude* remain unchanged between training (Train) and novel test set (Test). *Choice Consistency* and *Choice Accuracy* show a slight, expected decrease on the more challenging novel test set but remain high, confirming robust and effective generalization.

The networks demonstrated robust generalization, accurately selecting the higher-valued offer despite the stimulus novelty. Logistic regression confirmed that the core valuation strategy was preserved: key behavioral parameters, including inferred relative value and risk attitude, remained remarkably stable between constrained training and unconstrained test sets (Fig. 8C). While choice consistency and accuracy showed a slight, expected decrease on the novel set, overall performance remained high, confirming robust behavioral generalization.

This powerful generalization results from the network learning the task’s underlying mathematical structure, providing conclusive evidence for the multiplicative strategy sought earlier. The network learned a computation functionally equivalent to multiplication; a simpler, non-multiplicative strategy (e.g., additive heuristic) would lack a principle for evaluating these novel quantity-probability conjunctions, failing this test.

Crucially, this behavioral generalization was supported by a consistent neural implementation. We performed principal component analysis on the population activity of the networks as they processed novel offers. The analysis revealed that the underlying low-dimensional geometry of the decision remained unchanged. Population activity again separated into distinct, choice-specific clusters, with activity within each cluster graded by chosen value, mirroring the structure observed earlier (Fig. 4B; Supplementary Fig. S16). This shows the network applies the same low-dimensional computational strategy to familiar and novel situations, providing a robust neural basis for generalizable behavior.

Our findings thus align with and mechanistically explain observations in non-human primates, where subjects also generalize valuation processes to novel stimuli^22^. The model demonstrates how neural circuits can leverage fundamental computational principles, such as approximate multiplication, to achieve flexible, adaptive behavior extending beyond their direct experience.

## DISCUSSION

We developed a biologically plausible E/I RNN to advance and test hypotheses about the circuit basis of economic choice. Our primary contributions lie not in replicating known neural patterns, but in proposing concrete circuit mechanisms for fundamental computations. We show how: (1) a neural circuit with additive synapses can approximate non-linear multiplication of reward features, explaining generalization of learned values to unseen offers; (2) subjective economic preferences are physically stored in synaptic weights of long-range inputs to the decision circuit; (3) value comparison is implemented by a specific CRI motif, a detailed E/I connectivity pattern underlying WTA dynamics; and (4) a single circuit can achieve multitasking through compositional representations that flexibly allocate shared and specialized neural resources. Together, these findings constitute a coherent, mechanistically grounded framework for value-based decision making.

Although they were not explicitly trained for risk preference, our networks developed idiosyncratic risk attitudes, with some becoming mildly risk-averse and others risk-seeking (Figure 2D). This variability, likely arising from stochastic factors (e.g., weight initialization, trial history), mirrors individual differences observed in biological agents and suggests that such biases can arise naturally from reinforcement learning principles without a specialized objective. This provides a computational platform for testing hypotheses about the circuit origins of choice biases^33^, including those observed in neuropsychiatric disorders. For example, one could manipulate circuit properties (e.g., E/I balance) and examine consequences for risk preference and choice consistency.

This framework generates new, testable predictions. For instance, to induce risk aversion, one could train networks on skewed reward distributions. We predict a learned risk-averse policy would be implemented upstream of the decision circuit, through systematic changes in the input weights that distort the representation of probability, a mechanism consistent with neurophysiological findings in OFC^7,34^.

A central finding is that a model constrained by core biological principles reproduces a wide range of neurophysiological phenomena in the primate OFC. Neurons in the model exhibit tuning for key variables including offer value, chosen value, and choice^1,6^, with both excitatory and inhibitory cells displaying positive and negative tuning profiles. This heterogeneity, particularly the emergence of negatively tuned neurons, is a non-trivial, learned property of the recurrent circuitry. We propose that these neurons are not epiphenomenal but computationally critical for robust decisions, and that they enhance contrast between competing options. Targeted silencing of negatively tuned neurons impaired choice accuracy. This causally demonstrates their functional importance and offers a testable prediction for future optogenetic studies.

That both excitatory and inhibitory cells encode decision variables highlights a broader conceptual point about circuit computation. Modern experimental tools allow targeting specific cell types, which can encourage hypotheses like “cell type A performs function X”. In contrast, our work underscores that neural function is an emergent property of network interactions, not an isolated property of a cell class. For example, inhibitory neurons in our model are not simply ‘choice neurons’ or ‘offer value neurons’; rather, their structured connectivity within the recurrent E/I circuit *enables* the network to perform value comparison and generates the tuning required for robust computation. This network-centric perspective aligns with a long-standing tradition in computational neuroscience^23^ and provides a theoretical lens to interpret cell-type-specific data in the context of distributed computations.

At the population level, our results show that value computation and value comparison are organized within low-dimensional dynamics where specific neural activity patterns correspond to decision variables^32^, with chosen values potentially encoded as line attractors^35^. Our study provides testable hypotheses about the implementation of these computations. We propose that non-linear multiplication of reward features occurs upstream of the recurrent circuit and that learned relative values that define economic indifference points are physically stored in synaptic efficacy of these input projections. This architecture enables generalization to novel offers^22^ and explains how value comparison can be implemented by a competitive dynamics within the recurrent circuit itself^23,24,29^.

With respect to compositionality, our networks developed both shared and specialized representations to solve five distinct economic choice tasks. While four tasks elicited highly overlapping neural activity patterns, the sequential task required a specialized neural subspace, likely to support its unique working memory demands. In other words, our networks reused a core set of computational components for common operations (e.g., value integration and comparison) while flexibly engaging specialized modules for novel computational demands^25,26^. This flexi-ble resource allocation, facilitating accelerated learning via curriculum protocols^36,37^, suggests a general principle for how the brain balances task-specific adaptation with the efficiency of a shared architecture^38,39^.

Although motivated by experimental work in OFC, we intend our model to serve as a general framework for how a biologically plausible circuit can implement core computations for economic choice. This naturally leads to considering how this framework fits within a multi-regional architecture and engages with key debates in the field. An important example is the “Rudebeck/Murray challenge”, which posits a functional double dissociation between OFC and ventrolateral prefrontal cortex (VLPFC) based on lesion studies^11^. While neurophysiological work provides evidence that OFC neurons robustly encode economic variables including risk and probability^40–43^, our model does not presuppose that OFC is the sole locus of computation. As technical advances enable simultaneous recording from multiple brain regions, it is increasingly recognized that neural coding is distributed; value computation likely involves several cortical and subcortical regions. A major question is how to understand the functional specialization of brain regions like OFC compatible with distributed neural representations^44^. In fact, our framework is readily extendable to test distributed architectures. For instance, the abstract ‘probability’ input to our network could be modeled as an output from a VLPFC-like area, and their interaction could be studied. Future multi-regional models that can reconcile findings from both lesion and neurophysiology studies are a crucial next step, and our work provides a robust single-module component for such theories.

Our model presents a few limitations. Using an explicit task cue aligns with experimental paradigms^34^, but our analysis reveals that cues become largely redundant for four of the five tasks as a shared internal context is formed (Figure 7A). Similarly, modeling a fixation-maintenance signal at the same output level as good-based choices is a deliberate simplification. We acknowledge that our model is a framework for economic choice computation, not strictly a model of OFC anatomy. Indeed, as with virtually all trained RNNs to date, our model is limited to a single neural population, whereas the brain is a multi-regional system. In the brain, action selection and fixation control are likely managed by a broader network, including fixation-related neurons in the frontal eye fields (FEF) and superior colliculus (SC)^45^ and downstream areas like the lateral prefrontal cortex (LPFC), which must integrate value signals from OFC-like areas to guide behavior^46,47^. Investigating these multi-regional interactions, as well as uncued task switching^27^, represents a critical direction for future work.

In conclusion, our work establishes a robust computational framework for exploring the neural circuits underlying economic decisions. By providing a mechanistically-grounded and biologically-constrained model, we bridge single-neuron observations with population-level dynamics and offer a new suite of testable hypotheses to guide future empirical research into the neural basis of decision-making.

## RESOURCE AVAILABILITY

### Lead contact

Requests for further information and resources should be directed to and will be fulfilled by the lead contact, Xiao-Jing Wang.

### Materials availability

This study did not generate new unique reagents or biological materials.

### Data and code availability

All original code to train network models and generate data for this study have been deposited on GitHub at https://github.com/aldobattista/neural-circuit-economic-choice and are publicly available as of the date of publication. Any additional information is available from the lead contact upon reasonable request.

## ACKNOWLEDGMENTS

We thank G. Tavoni, C. Constantinople, E. Rich, and V. Fascianelli for their critical comments on the manuscript, as well as current and former members of the Wang lab, especially W. Soo, G. Benigno, Maryada, Y. Liu, V. Goudar, and A. Fanthomme, for fruitful discussions. This work was supported by the Swartz Foundation (A.B.); ONR grant N00014-23-1-2040 (X.-J.W.); NIH grants R01MH104494 (X.-J.W.), R01DA032758 (X.-J.W.), the National Institute of Neurological Disorders and Stroke (NINDS) and National Institute on Drug Abuse (NIDA) of the National Institutes of Health grant number U19NS123714 (X.-J.W.), and R01DA055709 (C.P.-S.); and the NYU IT High Performance Computing resources, services, and staff expertise.

## AUTHOR CONTRIBUTIONS

A.B., C.P.-S., and X.-J.W. conceptualized the study. A.B. developed the software, trained the networks, performed the data analysis, and visualized the results under the supervision of X.-J.W. C.P.-S. provided critical intellectual input throughout the project. A.B. wrote the initial draft of the manuscript. All authors contributed to manuscript revision and approved the final version.

## DECLARATION OF GENERATIVE AI AND AI-ASSISTED TECH-NOLOGIES

During the preparation of this work, the author(s) used Gemini 2.5 Pro (Google) to assist in improving the clarity and prose of the manuscript. After using this tool, the author(s) reviewed and edited the content as needed and take full responsibility for the content of the publication.

## SUPPLEMENTAL INFORMATION INDEX

Supplemental Information includes 16 figures and 2 videos and can be found with this article online.

**Figures S1–S16**. A single PDF document containing all supplementary figures, their legends, and descriptions for Supplementary Videos S1 and S2.

**Video S1**. Dynamic visualization of decision formation in an intact trained network, related to Figure 4. (MP4)

**Video S2**. Dynamic visualization of failed decision formation in a lesioned network, related to Figure S7. (MP4)

## MAIN FIGURES WITH TITLES AND LEGENDS

## STAR Methods

**Key resources table**

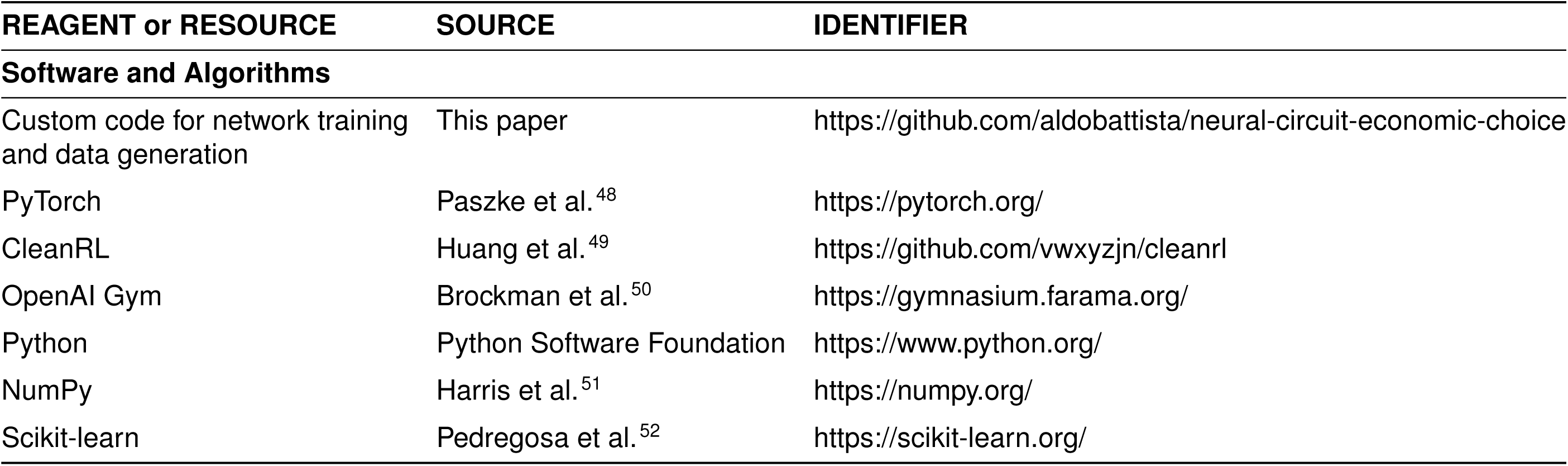

## Method details

### Network Model

#### Architecture and Dynamics

The model was an excitatory-inhibitory (E/I) continuous-time vanilla recurrent neural network (RNN)^18,25^, adhering to key biological constraints (Figure 1B). For the main multitasking analysis, we trained 20 independent networks. The networks consisted of *N* = 256 neurons, with an 80% excitatory and 20% inhibitory neuron ratio, reflecting cortical distributions^53^. Neurons had a single-unit time constant *τ* = 100 ms, consistent with NMDA receptor-mediated synaptic dynamics^23^. The dynamics of the network’s firing rate vector, **r**(*t*), were governed by:

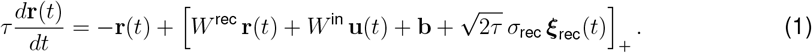

In this equation, *W*^rec^ is the recurrent weight matrix, *W*^in^ is the input weight matrix, **u**(*t*) is the input vector, and **b** is a bias term. The term ***ξ***_rec_(*t*) represents Gaussian white noise with a standard deviation of *σ*_rec_ = 0.15. The network’s dynamics were integrated using the Euler method with a temporal discretization of *δt* = 20 ms. The rectified linear unit (ReLU) activation function, [·]_+_, ensured non-negative firing rates.

The input vector **u**(*t*) included scalar representations of a fixation signal, the quantities and probabilities of offered goods, and task-specific rule cues. All scalar inputs were normalized between 0 and 1 and included a baseline of *u*_0_ = 0.2 and additive input noise:

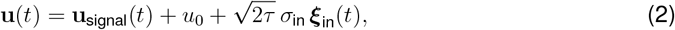

where **u**_signal_(*t*) represents the normalized input signals and ***ξ***_in_(*t*) is Gaussian white noise with a standard deviation of *σ*_in_ = 0.01.

#### Biological Constraints and Readouts

The network architecture strictly adhered to Dale’s Law: outgoing weights from excitatory neurons were constrained to be non-negative, and those from inhibitory neurons were non-positive. These constraints were enforced by applying fixed masks to the weight matrix during training^18^. Similarly, all long-range projections (input and output weights) were constrained to be excitatory^54^.

The network produced two distinct outputs from separate linear readouts: a policy for action selection (the “actor”) and a value function predicting the expected future reward (the “critic”). Crucially, both readouts originated from the same recurrent network, a design choice that aligns with experimental evidence that value-related areas are causally involved in choice^9^ and avoids the biological implausibility of entirely separate actor-critic modules^17^. The action space available to the actor included selecting one of the available offers or maintaining fixation, treating fixation as a distinct “hold” or “no-go” action. This integrated action space allows the network to learn not only what to choose but also when to execute the choice.

#### Initialization of Parameters

All weight matrices and biases were initialized as follows:

1. **Recurrent weights** (*W* ^rec^) were sampled from a Gamma distribution (shape and scale parameters of 4) and then rescaled so the spectral radius of the matrix was 1.5. The Dale’s Law mask was then applied.
2. **Input weights** (*W* ^in^) were sampled from a uniform distribution over 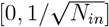, where *N*_*in*_ = 16 is the number of inputs.
3. **Output weights** 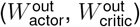 were sampled from a uniform distribution over 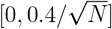.
4. **All biases** (**b, b**_actor_, *b*_critic_) were initialized to zero.

### Task Design and Training Paradigms

Networks were trained on a diverse set of five economic choice tasks designed to simulate aspects of primate decision-making experiments^7^. Each task required the network to evaluate and compare offers by integrating information about different goods, quantities, probabilities, and temporal structures. To investigate different aspects of learning and representation, networks were trained in three distinct paradigms: (1) *multitasking*, where a single network was trained concurrently on all five tasks; (2) *single-task training*, where a separate network was trained for each individual task; and (3) *curriculum learning*, where a single network was trained sequentially on tasks of increasing complexity.

#### Common Task Structure and Stimuli

Each trial began with a fixation period of variable duration drawn from a uniform distribution between 500 and 1500 ms, followed by a rule cue (uniformly distributed, 500 − 1500 ms) indicating the current task. After offer presentation, a response period of up to 1000 ms was provided for the network to make a choice. The variable epoch durations prevent the network from relying on temporal cues and encourage it to learn stable, state-dependent computational policies (e.g., attractor states) (Figure 1A).

#### Network Inputs and Rewards

While tasks were visualized with colored circles (Figure 1A), the network itself received scalar inputs. Dedicated input channels encoded the quantity and probability for each offered good. The intrinsic values of the goods (e.g., *ρ*_*A*_ = 3, *ρ*_*E*_ = 1) were not provided as inputs but were learned by the network from reward feedback based on choice outcomes. The fixation and task rule cues were also provided as distinct scalar inputs, representing abstract contextual and task-set information. Prematurely breaking fixation resulted in trial abortion and a negative reward of −1. Choosing an offer yielded a terminal reward. For deterministic offers, the reward was equal to the offer’s value (*ρ*_*X*_ *×* Quantity_*X*_, for good *X*). For probabilistic offers, a reward of (*ρ*_*X*_ *×* Quantity_*X*_) was delivered with probability *p*_*X*_, and a reward of 0 was delivered with probability (1 − *p*_*X*_).

#### Offer Value Calculation

The value of each offer was calculated as:

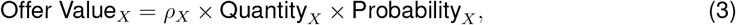

where *ρ*_*X*_ is the intrinsic value of the good *X*, Quantity_*X*_ was sampled uniformly from a range of 0 to 10*/ρ*_*X*_ (to ensure comparable value ranges across goods), and Probability_*X*_ was sampled uniformly from 0 to 1. In deterministic tasks, probability was set to 1. Finally, to ensure all choices presented to the network were non-trivial, a threshold was applied: if the computed Offer Value for a given offer was less than 1, its quantity and probability parameters were re-sampled until the criterion was met.

#### Task Details

The five tasks were designed to impose distinct computational demands:

- **Standard Task:** Choice between two goods (C vs. E) varying only in quantity. Demands basic value comparison.
- **Risky Task:** Choice between two goods (C vs. E) varying in both quantity and probability. Requires multiplicative integration of features.
- **Bundles Task:** Choice between two bundles of goods (e.g., B+C vs. D+E). The total value of each bundle was the sum of its components’ expected values. Requires summation of values within an offer.
- **Ternary Task:** Three-way choice between goods A, C, and E. Increases the dimensionality of the choice space.
- **Sequential Task:** Choice between two goods (C vs. E) presented sequentially with an intervening delay. Requires working memory to hold the value of the first offer for comparison.

### Reinforcement Learning Procedure

#### Algorithm and Loss Function

We trained the networks using Proximal Policy Optimization (PPO)^20,49^, a state-of-the-art policy gradient algorithm that optimizes the policy directly to maximize the expected cumulative reward. The total loss function ℒ (*θ*) to be maximized is a weighted sum of a policy loss, a value function loss, and an entropy bonus to encourage exploration:

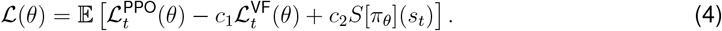

The coefficients *c*_1_ = 0.5 and *c*_2_ = 0.01 are hyperparameters. The expectation E is taken over batches of experience collected by running 20 parallel environments for *T* = 128 steps, where each step corresponds to one simulation time step of *δt* = 20 ms.

The PPO policy loss uses a clipped objective function to ensure stable policy updates:

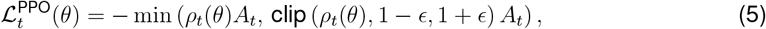

where 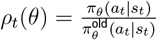 is the probability ratio between the current and old policies, and *ϵ* = 0.1 is the clipping parameter.

The policy update is guided by the advantage function *A*_*t*_, which is analogous to the reward prediction error (RPE) in neuroscience^55,56^. It is calculated as the difference between the empirical return and the value function’s prediction:

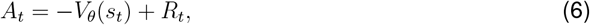

where *V*_*θ*_(*s*_*t*_) is the value function’s estimate for state *s*_*t*_, and *R*_*t*_ is the bootstrapped return, computed as:

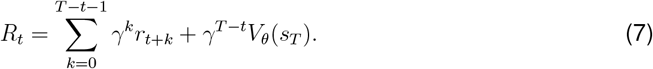

Here, *r*_*t*_ is the reward at time step *t*, and *γ* = 0.99 is the discount factor.

The value function loss trains the critic to accurately predict the bootstrapped return:

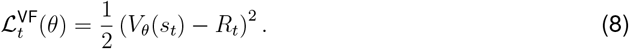

The entropy bonus encourages exploration:

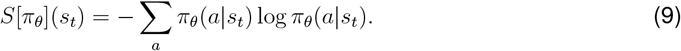

#### Optimization and Performance Criteria

All trainable parameters (*θ*) were updated via gradient ascent and backpropagation through time using the Adam optimizer with a learning rate of 2.5 *×* 10^−4 57^. The gradient norm was clipped at 1.0 to prevent exploding gradients. For curriculum learning protocols, networks were trained on simpler tasks before progressing to more complex ones. For example, a network might first be trained on the standard task with two deterministic options, then introduced to probabilistic outcomes in the risky task, and finally trained on the ternary task involving a third choice option.

Training continued until the network met two stringent performance criteria on a held-out test set of 25, 000 trials (∼ 5, 000 per task): (1) completing at least 99% of trials without breaking fixation, and (2) achieving at least 90% accuracy on correctly completed trials (Figure S1). The learning curves show that networks typically progressed through three distinct phases: first, learning to maintain fixation throughout the trial without making a choice; second, learning to select an action randomly during the response period; and finally, learning the value-based policy of selecting the highest-value offer (Figure S1C). All code was implemented in PyTorch^48^, Gym^50^, NumPy^51^, and Scikit-learn^52^.

### Quantification and statistical analysis

#### Behavioral Analysis via Logistic Regression

To quantitatively characterize network decision-making and extract behavioral parameters, we performed logistic regression analyses on the choice data from each task, following methodologies applied in primate studies^7^.

#### Data Collection

For each network, we collected 25, 000 trials (about 5, 000 trials per task) where the network made choices among different offers. Each offer was defined by its quantity (*q*_*X*_), probability (*p*_*X*_) when applicable, and ground-truth intrinsic value (*ρ*_*X*_). The intrinsic values were set during training, with *ρ*_*A*_ *> ρ*_*B*_ *> ρ*_*C*_ *> ρ*_*D*_ *> ρ*_*E*_.

#### Models for Binary Choices

In tasks involving choices between two options—the *standard, risky, bundles*, and *sequential* tasks—we modeled the probability of choosing one option over another using logistic regression. The models estimate a set of free parameters 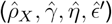 that best explain the network’s choice behavior. The estimated relative value, 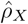, is defined relative to a reference good (good E), for which we set 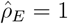.

#### Standard Task

In the *standard task*, the probability of choosing good C over good E is modeled as:

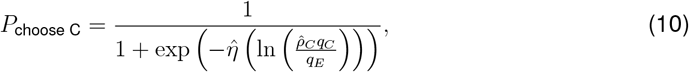

where 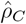 and 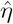 are the estimated relative value and choice consistency parameters, respectively.

#### Risky Task

In the *risky task*, we also estimate the risk attitude parameter, 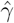:

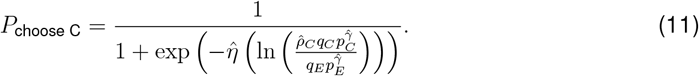

#### Bundles Task

In the *bundles task*, the probability of choosing Bundle 1 (B+C) over Bundle 2 (D+E) is modeled as:

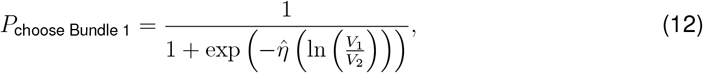

where 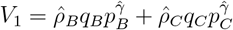 and 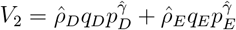 (with 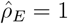).

#### Sequential Task

In the *sequential task*, we additionally estimate the order bias parameter,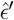

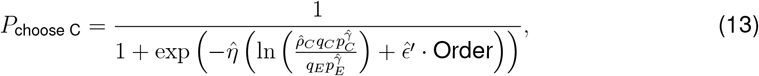

where Order = +1 if good C was presented second, and −1 if it was presented first.

#### Model for Multinomial Choices

In the *ternary task* (choices among A, C, E), we used multinomial logistic regression:

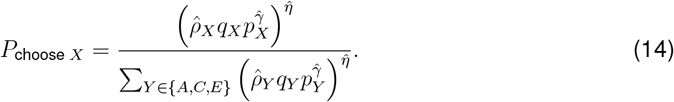

#### Parameter Estimation

We estimated all behavioral parameters 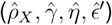 using maximum likelihood estimation for each task and network individually. The estimated relative values, denoted by 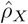 to distinguish them from the ground-truth values *ρ*_*X*_ used in training, are relative measures, with 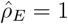 serving as the reference point, allowing for comparison across tasks and networks. The choice consistency parameter 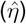 indicates the steepness of the psychometric function; higher values correspond to more consistent choices based on offer value differences.

### Single-Neuron Analyses

#### Data Collection and Trial Structure

For neural analyses, we collected test trials (25, 000 per network) with fixed epoch durations to facilitate temporal alignment of activity across trials. In the sequential task, neural activity was aligned to the onset of the *second* stimulus, as this is the point where all information required for the decision becomes available to the network.

#### Single-Neuron Selectivity Analysis

To analyze neuronal selectivity for decision variables, we performed linear regression of each neuron’s firing rate (*r*(*t*)) against each candidate behavioral variable (*X*) independently at each time point: *r*(*t*) = *β*_0_ + *βX* + *ϵ*. The variables considered included offer values (OVC, OVE), chosen value (CV) and choice (CH). These analyses were performed on a set of approximately 5,000 test trials per task for each of the 20 networks.

To determine whether a neuron was selective for a given variable at a specific time point, we applied a standard two-step statistical criterion. First, the linear regression had to be statistically significant (*p <* 0.05). Second, to ensure a minimum effect size, the regression had to explain a baseline amount of variance (*R*^2^ ≥ 0.005). For analyses where a neuron was assigned to a single best-fit variable category (e.g., for the fraction of selective neurons plot in Figure 3C), we chose the variable that yielded the maximum *R*^2^ among all variables that satisfied both criteria. If no variable met these criteria, the neuron was considered non-selective at that time point.

#### Temporal Stability Index (TSI)

For each neuron deemed selective at any point during the trial, its “primary variable” was identified as the one it encoded for the majority of time steps. The TSI was then calculated as:

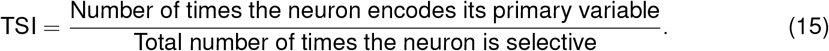

A TSI of 1 indicates that the neuron consistently encodes the same variable whenever it is selective, while lower values suggest dynamic selectivity, where a neuron’s tuning changes over time.

#### Analysis of Categorical and Mixed Selectivity

To investigate the nature of neural encoding, we analyzed the distribution of differences in the coefficient of determination 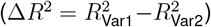 for pairs of decision variables. This approach was chosen to directly parallel analyses in primate neurophysiology^28^. We performed this analysis using two different criteria for neuronal inclusion to distinguish between the coding schemes of specialized versus general neuronal populations. **Categorical encoding in highly selective neurons (Main Text, Figure 3E):** To test for categorical encoding within the most specialized neurons, we applied a stringent inclusion threshold. For each pair of variables, we included a neuron-time point in the analysis only if it was very strongly modulated by at least one of the two variables (*R*^2^ ≥ 0.5). The resulting Δ*R*^2^ distributions for this subpopulation were plotted to assess their modality.

#### Mixed selectivity in the general population (Supplementary Figure S4)

To characterize the coding scheme of the broader population of task-related neurons, we used a more lenient inclusion criterion. Here, a neuron-time point was included if it was significantly modulated by at least one of the variables (*p <* 0.05 and *R*^2^ ≥ 0.005). The Δ*R*^2^ distributions for this general population were then analyzed. Together, these two analyses provide a comprehensive view of the network’s dual coding strategy.

### Neural Population Analyses

#### Population State-Space Analysis and Dimensionality

To investigate population-level encoding, we performed principal component analysis (PCA) on the activity of the *N* = 256 recurrent neurons. For each task, the data matrix for PCA was constructed from the neural activity during the last 200 ms of the stimulus presentation phase, concatenated across all available test trials (approx. 5, 000 per task). This resulted in a matrix of size *N ×* (num trials *×* num time steps). We analyzed the full population as well as the excitatory and inhibitory populations separately.

We estimated the effective dimensionality of the neural activity using the participation ratio (PR). The PR is a standardized, continuous measure of dimensionality derived from the eigenvalues (*λ*_*i*_) of the activity’s covariance matrix, which avoids the need for an arbitrary variance threshold. Intuitively, it quantifies how broadly the variance of population activity is distributed across principal components. If all variance is concentrated in a single dimension, the PR is 1; if variance is spread evenly across *M* dimensions, the PR is *M* . It is formally defined as:

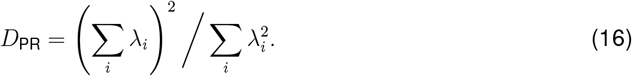

#### Principal Component Regression

To interpret the dimensions of population activity, we projected the single-trial neural activity onto the first three principal components. We then performed a multiple linear regression where the dependent variables were the projections onto each PC, and the independent variables were the key decision variables for that trial (e.g., offer values, chosen value, choice). The coefficient of determination (*R*^2^) from these regressions quantifies the proportion of variance in each principal component that is explained by each decision variable.

#### Computational Lesion Analyses

To causally dissect the contributions of different circuit elements, we performed computational lesion experiments after network training was complete. The primary lesion reported here involved removing all recurrent connections by setting the recurrent weight matrix to zero (*W* ^rec^ = 0), thereby isolating the feedforward processing pathway. After this lesion, we re-evaluated the network on a full set of test trials and analyzed two key outcomes: (1) Behavioral Performance, by quantifying the performance drop in trial completion rate (Figure S6B), and (2) Neural Activity, by re-calculating firing rate distributions and repeating the full PCA and dimensionality analyses on the lesioned network activity (Supplementary Figures S6A and S7). This allowed for a direct comparison of the population geometry and behavioral capacity between the intact and lesioned circuits.

#### Visualization of Population Dynamics

To visualize the evolution of the population state, we created both static plots and dynamic videos. For static plots like Figure 4B, we show the state of all trials projected onto the first two PCs at a single, representative time point (the end of the offer period). For dynamic visualizations (Supplementary Videos S1 and S2), we created an animation by plotting this full cloud of trial-points at each successive 20 ms time step over the course of the entire trial. This method allows visualization of how the entire distribution of population states evolves during decision formation.

### Analyses of Input Weights and Upstream Computation

#### Analysis of Input Weight Structure and Mixed Selectivity

To directly test for the existence of mixed-selective neurons hypothesized to cause the geometric rotation of the value representation, we analyzed the input weight matrices (*W*^in^) of trained networks. We first visualized the overall structure by computing the Pearson correlation coefficients between the input weight vectors for all offer features (Fig. 5A). To provide a quantitative and direct measure of input integration, we calculated a Selectivity Index (SI) for each recurrent neuron. This analysis was performed on the 50 networks trained exclusively on the risky task (the same cohort used for the analysis in Fig. 5E). For each recurrent neuron *i* in each network, the SI was calculated based on its synaptic input weights from the quantity of good C (*w*_*qC*_) and good E (*w*_*qE*_) as follows:

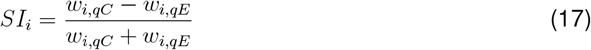

The SI was defined using only quantity weights because the input weights for the quantity and probability of any given offer were found to be highly correlated (Fig. 5A), making the quantity-based index a robust measure of a neuron’s overall offer preference. An SI value of +1 or −1 indicates a neuron is exclusively driven by input from good C or good E, respectively, while an SI of 0 indicates perfectly balanced inputs from both. For each of the 50 networks, we computed the probability density function of the SI values across its recurrent population. These 50 distributions were then averaged to produce the final histogram shown in Supplementary Figure S9, which visualizes the converged wiring strategy found by the learning algorithm.

#### Testing for Multiplicative Computation in the Feedforward Pathway

To investigate the circuit mechanism responsible for the emergent multiplicative-like computation, we performed a computational dissection to isolate the feedforward pathway from the recurrent circuitry. After training was complete, we analyzed networks with their recurrent connections computationally lesioned (e.g., setting the recurrent weight matrix *W* ^rec^ = 0). We collected neural activity from the last 200 ms of the stimulus presentation phase across approximately 5, 000 trials of the risky task. We then performed Principal Component Analysis (PCA) on the resulting feedforward population activity. To determine if this isolated pathway supported a multiplicative or additive integration scheme, we used linear regression to compare how well two different models could explain the variance in the top two principal components (PCs). The first model used the multiplicative product of quantity and probability for each offer (OVC and OVE) as predictors. The second model used their additive sum (OVCs and OVEs). Higher coefficients of determination (*R*^2^) for the product-based model were taken as strong evidence that the feedforward architecture had learned to perform a non-linear, multiplicative computation (Fig. 5B).

#### Toy Feedforward Network Model for Multiplication and Mixed Selectivity

To provide a proof-of-principle for how a neural circuit can both approximate multiplication and generate a rotated value representation, we constructed a simplified feedforward network. The demonstration is two-fold: first, we provide a constructive proof that the architecture can approximate a multiplicative function, and second, we use this capability to test the mechanistic origin of the geometric rotation observed in the full network.

##### Constructive Proof of Multiplicative Approximation

Our proof rests on the principle of function approximation with basis functions. The goal is to approximate the non-linear value surface *V* (*q, p*) = *q* · *p* by forming a linear combination of simpler, non-linear basis functions provided by a population of neurons. The activity *r*_*i*_ of a single neuron *i* with a ReLU activation provides a simple, ramp-like basis function:

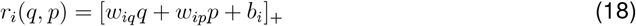

where *q* and *p* are inputs, *w*_*iq*_ and *w*_*ip*_ are weights, and *b*_*i*_ is a bias. A single such neuron is insufficient. However, a population of *N* neurons, each with different weights and biases, can generate a diverse set of these basis functions that tile the input space. A downstream linear readout can then learn to weight the contribution of each neuron to reconstruct the target multiplicative surface. We provide a constructive proof of this by engineering the weights and biases to achieve this tiling systematically. For the *i*-th neuron in a population of size *N*, we set the parameters as:

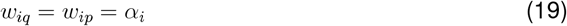

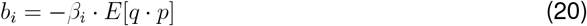

where *E*[*q* · *p*] is the mean value of the product over the task’s offer distribution. By making *α*_*i*_ and *β*_*i*_ vary across the population (e.g., linearly with neuron index *i*), we ensure that different neurons become active at different locations and with different sensitivities across the quantity-probability space. This architecture is thus mathematically capable of approximating multiplication. We provide empirical validation for this constructive proof with a parameter search (Supplementary Fig. S8). In this search, where for simplicity all neurons shared the same *α* and *β*, we demonstrated that there exists an optimal parameter regime where the population activity is overwhelmingly better explained by a multiplicative (*q* · *p*) than an additive (*q* + *p*) model. This confirms that the architecture is not only capable but also predisposed to learn a multiplicative integration rule. *Modeling Mixed Selectivity to Explain Geometric Rotation:* Having established the network’s ability to compute value, we applied this module to test the origin of the value sum/difference representation.

1. *Segregated Model* (Fig. 5C): We used two independent populations, each approximating the value of one offer, to show this wiring is insufficient to produce the observed rotation.
2. *Mixed-Selectivity Model* (Fig. 5D): We added a third population receiving summed inputs from both offers. This structure, mimicking our analysis of the full network’s weights (Supplementary Fig. S9), successfully reproduced the geometric rotation, thereby providing a causal, mechanistic explanation for this emergent geometric feature.

#### Correlation of Relative Values and Input Weights

To test the hypothesis that learned subjective values are physically encoded in the synaptic strengths of input projections, we performed a dedicated experiment designed to establish a causal link between a manipulated training parameter, the resulting circuit structure, and the emergent behavioral preference. We trained a total of 50 independent networks exclusively on the risky task, organized into five groups of ten. For each group, the intrinsic value of the reference good E was held constant (*ρ*_*E*_ = 1), while the intrinsic value of the target good C was systematically set to a different ground-truth value: *ρ*_*C*_ ∈ {1, 2, 3, 4, 5} . After each of the 50 networks was trained to the standard performance criteria, we extracted two key quantities. First, to capture the network’s learned preference, we inferred its behavioral relative value, 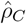, by fitting the logistic regression model for the risky task to 5, 000 of its choice trials. Second, to quantify the underlying circuit parameter, we calculated the average input synaptic strength by taking the mean of all non-zero excitatory weights connecting the input for the quantity of good C (*q*_*C*_) to the recurrent population. Finally, we performed a linear regression across the 50 resulting pairs of values (inferred behavioral value vs. average input weight) to quantify the relationship between the network’s learned behavior and its input circuit structure, as shown in Figure 5E.

### Analyses of Recurrent Circuitry and Value Comparison

#### Analysis of Output Dynamics

To characterize the dynamics of the decision process at the action-selection level, we analyzed the activity of the network’s output units during the risky task. These units include the policy outputs (the “actor”) corresponding to available actions (e.g., ‘Choose C’) and the value function output (the “critic”). For Figure 6A, we averaged the output activity across all trials, separating trials based on the network’s final choice. For Figure 6B, we constructed a heatmap summarizing a series of linear regressions. Each row of the heatmap corresponds to an output node, and each column to a decision variable. The *R*^2^ value in each cell represents the coefficient of determination from a simple linear regression of a single output node’s activity against a single decision variable, performed across all trials for a given task. To reveal choice-specific encoding, the set of regressors included conjunctive variables. For instance, the variable ‘CVBC’ (‘Chosen Value BC’) was defined as the value of good C or B+C on trials where C or B+C was chosen, and was set to zero on all other trials. The winner-take-all (WTA) dynamic is demonstrated by a choice-specific output (e.g., ‘BC Ch’) being strongly predicted by its corresponding conjunctive variable (e.g., ‘CVBC’) but not by the variable for the competing choice (e.g., ‘CVDE’).

#### Reaction Time Analysis

Reaction times (RTs) were measured from the onset of the response phase (when the fixation signal turned off) to the time step at which the network executed a choice action. We collected RTs across all correctly completed trials for all tasks. We plotted the probability density of these RTs and, to assess the relationship between decision difficulty and response latency, performed a linear regression between RTs and the absolute value difference of the two offers on a trial-by-trial basis (Supplementary Fig. S10A).

#### Recurrent Connectivity Analysis

To identify learned connectivity patterns, we examined the recurrent weight matrices (*W*^rec^) of the trained networks (Supplementary Fig. S10B, left). To assess the effective dimensionality of this connectivity, we performed a Singular Value Decomposition (SVD) on the full 256 *×* 256 weight matrix for each network. A low-rank matrix, indicated by a spectrum of singular values that decays rapidly, implies that the network’s dynamics are governed by a few dominant connectivity motifs (Supplementary Fig. S10B, right).

#### Reduced Connectivity Matrix and Circuit Diagram

To abstract the core functional architecture from the full connectivity matrix and make it interpretable, we performed a many-to-one functional abstraction, visualized for a single network in Supplementary Fig. S11. This procedure consists of three steps. First, for a given task (with the risky task used for the main analysis in Figure 6), we performed a regression analysis to classify each of the 256 individual neurons into a functional pool based on its cell type (excitatory or inhibitory) and its primary selectivity (e.g., ‘Choice C’). Second, all neurons with the same classification were grouped into a single functional pool. Third, we constructed a reduced connectivity matrix by computing the mean synaptic weight of all connections between each pair of neuron pools. The final matrix shown in Figure 6C represents the average of these reduced matrices across all 20 networks. This matrix highlights the dominant connectivity motifs, from which we created the simplified circuit diagram in Figure 6D. To test the generality of this finding, this entire procedure was repeated for the standard, bundles, ternary, and sequential tasks, revealing that the same core Competitive Recurrent Inhibition (CRI) motif emerged in all choice contexts (Supplementary Fig. S10C).

#### Lesion Experiments for Causal Validation

To causally test the functional roles of specific circuit elements, we performed two targeted computational lesion experiments on the fully trained networks.

1. *Lesion of Negatively Tuned Neurons:* To test the hypothesis that negatively tuned neurons are computationally critical, we first identified all neurons in each of the 20 multitask networks that had a statistically significant negative regression slope (*p <* 0.05) for either ‘Offer Value’ or ‘Chosen Value’ variables during the offer period. During subsequent test simulations across all five tasks, the activity of these specific neurons was clamped to zero. We then measured the percentage change in performance (both trial completion and choice accuracy) relative to the intact network (Supplementary Fig. S12).
2. *Functional Ablation of Neuronal Pools:* To identify the most critical functional populations, we performed a systematic ablation study on the 20 multitask networks, tested on the risky task. We first classified all neurons into functional pools as described above. We then, in separate experiments, silenced each functional pool by clamping the activity of all its member neurons to zero throughout the trial. We measured the percentage change in trial completion and choice accuracy for each ablation experiment to determine the causal importance of each cell group (Supplementary Fig. S13).

### Analyses of Compositionality and Learning

#### Analysis of Rule Cue and Neural Subspaces

To investigate how networks represent different tasks, we analyzed both the input weights and the resulting population activity of the 20 multitasking networks. First, we extracted the input weight vectors (*W* ^in^) connecting each of the five rule cue inputs to the recurrent neurons. We then computed the pairwise Pearson correlation between these vectors for each network and averaged the resulting correlation matrices across all 20 networks to produce the matrix shown in Fig. 7A. Second, for the subspace analysis (Fig. 7B), we performed principal component analysis (PCA) on the population activity during the last 200 ms of the rule cue period for each task, using data concatenated across all 20 networks. To determine the intrinsic dimensionality of each task’s representation, we first calculated the Participation Ratio (PR) for each task’s activity. The number of principal components (*n*_*PC*_) used to define the subspace for each task was then set to the closest integer to its calculated PR value. Finally, we computed the principal angle between the subspaces spanned by the top *n*_*PC*_ principal components for each pair of tasks. A smaller angle indicates more overlapping, or shared, neural representations.

#### Task Variance and Neuronal Clustering

To identify shared and specialized neural populations in multitasking networks, we calculated the variance of each neuron’s firing rate during the stimulus presentation phase across all trials for each task. This variance was then normalized for each neuron by its maximum variance across all five tasks, creating a 5-dimensional task-variance profile for each neuron. For each network, we then used the *k*-means clustering algorithm (from scikit-learn^52^) on these profiles to group neurons based on their functional contribution to different tasks. The analysis was performed independently on the excitatory and inhibitory populations. For each population in each network, the optimal number of clusters (*k*) was determined by maximizing the silhouette score across a range of possible *k* values (from 2 to 20). While the precise optimal number of clusters could vary slightly between networks, the qualitative functional organization was highly consistent. The heatmaps in Figure 7C show the results from a representative network, for which the optimal number of clusters was found to be *k* = 3 for the excitatory population and *k* = 2 for the inhibitory population. Critically, across all networks, this procedure consistently identified a ‘shared’ cluster (neurons active in all tasks) and ‘specialized’ clusters (neurons with high variance in only one or two tasks, such as the ternary and sequential tasks).

#### Lesion Analysis of Neuronal Clusters

To causally assess the functional importance of the identified neuronal clusters, we performed a series of targeted lesion analyses on the representative multitasking network shown in Figure 7C. For each of the excitatory and inhibitory clusters identified in this network (Shared, Ternary-specialized, and Sequential-specialized), we performed a separate lesion experiment. During test trials across all five tasks, we computationally silenced all neurons belonging to the targeted cluster by clamping their firing rates to zero throughout the entire trial. We then evaluated the network’s performance by measuring two metrics: (1) the Trial Completion Rate and (2) the Accuracy on successfully completed trials. The results, shown in Fig. S14, are presented as the percentage drop in performance for the lesioned network compared to its intact (unlesioned) control performance.

#### Fractional Task Variance Analysis

For each pair of tasks in the 20 multitasking networks, we computed the fractional variance difference for each neuron, defined as (*TV*_1_ − *TV*_2_)*/*(*TV*_1_ + *TV*_2_), where *TV*_1_ and *TV*_2_ are the normalized firing rate variances for the two tasks. We then plotted histograms of these values across the entire neural population (Fig. 7D) to visualize the organizational relationship between the neural resources recruited for different tasks.

#### Visualization of Task Representations in a Common PCA Space

To visualize the compositional structure across all 20 independently trained multitasking networks, it was necessary to align their individual state spaces. Because PCA yields a basis set whose component signs and order are arbitrary, the principal components are not directly comparable across networks. We therefore implemented a rotational alignment procedure using a Procrustes analysis. First, we designated one network as the reference. Then, for each of the other 19 networks, we found the optimal orthogonal rotation matrix that minimized the sum of squared distances between its mean task representations and those of the reference network. After applying this rotation to align each network’s state space to the common reference, we projected the mean population activity for each of the five tasks from all 20 networks into this single, aligned space to create the visualization in Fig. 7E.

#### Zero-Shot Transfer Learning Analysis

To directly test whether multitasking is necessary for developing a flexible, multi-skilled circuit, we performed a zero-shot transfer learning analysis (Fig. S15). We trained five separate groups of networks (10 networks per group), with each group trained exclusively on one of the five tasks until the standard performance criteria were met. After training on its single source task, each network was evaluated on a held-out test set of 5, 000 trials from all five tasks without any further weight updates. We calculated and plotted two performance metrics for this cross-task evaluation: the Trial Completion Rate and the Accuracy on successfully completed trials.

#### Curriculum Learning Protocols

To assess whether prior experience accelerates the acquisition of new skills, we implemented curriculum learning protocols (Fig. 7F). We focused on three complex target tasks: Ternary, Bundles, and Sequential. For each target task, we compared the number of training trials required to reach the performance criterion under three distinct training curricula. A separate group of 10 independent networks was trained for each condition, for a total of 90 networks in this analysis (3 target tasks *×* 3 curricula *×* 10 networks). The three curricula were:

1. *From Scratch* (Rand): Networks were trained on the target task directly from a random weight initialization.
2. *Single Pre-training* (Rand→Stand): Networks were first trained to criterion on the Standard task, and then training continued on the target task.
3. *Double Pre-training* (Rand→Stand→Risky): Networks were first trained to criterion on the Standard task, then to criterion on the Risky task, and finally trained on the target task.

The box plots in Fig. 7F show the number of training trials required to learn only the *final* target task in each curriculum. To facilitate direct comparison on a common scale, for each target task (e.g., Ternary), the trial counts for all three curriculum conditions were normalized by the maximum number of trials required by any single network across all conditions for that specific task. This ensures that the y-axis in each panel ranges from 0 to 1, representing the fraction of the maximum observed learning time.

### Analyses of Generalization to Novel Offers

#### Generalization Test Design

To test if the network could generalize a learned valuation rule to novel stimuli, we trained ten networks exclusively on the risky task using a constrained, orthogonal set of offers. During this *training phase*, for each trial, offers for goods C and E were generated under one of two conditions, chosen randomly with 50% probability: (1) probabilities for both goods were fixed at 0.5 while quantities varied across their full range, or (2) quantities were fixed at half their maximum value (e.g., *q*_*C*_ = 2.5, *q*_*E*_ = 5) while probabilities varied across their full range (0 to 1). This orthogonal design (visualized in Fig. 8A) ensured that the network never encountered offers where both quantity and probability varied simultaneously from their central values, forcing it to learn their contributions to value independently. Training continued until networks achieved high performance (≥ 90% accuracy) on this constrained set. During the subsequent *testing phase*, we assessed generalization by evaluating the fully trained networks on a new, unconstrained set of 5, 000 offers where both quantity and probability for both goods varied independently and simultaneously across their full ranges. This test set was therefore composed almost entirely of novel quantity-probability combinations that the networks had not encountered during training.

#### Behavioral Analysis

We performed logistic regression analysis on the choice data from both the constrained training set and the unconstrained test set for each of the ten networks. We used the same logistic model defined for the risky task in the “Behavioral Analysis via Logistic Regression” section. By fitting this model separately to the training and test data, we extracted and compared the key behavioral parameters (inferred relative value 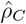, risk attitude 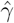, choice consistency 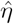, and choice accuracy) to assess the consistency of the network’s learned valuation strategy when faced with novel offer combinations (Fig. 8C).

#### Analysis of Neural Dynamics during Generalization

To determine if the neural implementation of the decision generalized along with the behavior, we analyzed the population activity of the ten constrained-trained networks during the test phase with novel, unconstrained offers. We collected neural activity from the last 200 ms of the stimulus presentation phase across all test trials (5, 000 trials per network). We then performed principal component analysis (PCA) on the population activity to visualize the structure of the neural state space, as shown for a representative network in Supplementary Fig. S16. The consistency of the PCA geometry (Supplementary Fig. S16) with that of the standard multitasking networks (Fig. 4B) demonstrates that the same low-dimensional computational solution was employed for both familiar and novel stimuli.

## References

1. Padoa-Schioppa, C. (2011). Neurobiology of economic choice: a good-based model. Annual review of neuroscience 34, 333–359.

2. Padoa-Schioppa, C., and Conen, K.E. (2017). Orbitofrontal cortex: a neural circuit for economic decisions. Neuron 96, 736–754.

3. Platt, M.L., and Glimcher, P.W. (1999). Neural correlates of decision variables in parietal cortex. Nature. 400, 233–238.

4. Sugrue, L.P., Corrado, G.C., and Newsome, W.T. (2004). Matching behavior and representation of value in parietal cortex. Science 304, 1782–1787.

5. Barraclough, D.J., Conroy, M.L., and Lee, D. (2004). Prefrontal cortex and decision making in a mixed-strategy game. Nat. Neuroscience 7, 404–410.

6. Padoa-Schioppa, C., and Assad, J.A. (2006). Neurons in the orbitofrontal cortex encode economic value. Nature 441, 223–226.

7. Padoa-Schioppa, C. (2022). Logistic analysis of choice data: A primer. Neuron 110, 1615– 1630.

8. Padoa-Schioppa, C., and Assad, J.A. (2008). The representation of economic value in the orbitofrontal cortex is invariant for changes of menu. Nature neuroscience 11, 95–102.

9. Ballesta, S., Shi, W., Conen, K.E., and Padoa-Schioppa, C. (2020). Values encoded in orbitofrontal cortex are causally related to economic choices. Nature 588, 450–453.

10. Ballesta, S., Shi, W., and Padoa-Schioppa, C. (2022). Orbitofrontal cortex contributes to the comparison of values underlying economic choices. Nature communications 13, 4405.

11. Rudebeck, P.H., Saunders, R.C., Lundgren, D.A., and Murray, E.A. (2017). Specialized representations of value in the orbital and ventrolateral prefrontal cortex: desirability versus availability of outcomes. Neuron 95, 1208–1220.

12. Raghuraman, A.P., and Padoa-Schioppa, C. (2014). Integration of multiple determinants in the neuronal computation of economic values. Journal of Neuroscience 34, 11583–11603.

13. Solway, A., and Botvinick, M.M. (2012). Goal-directed decision making as probabilistic inference: a computational framework and potential neural correlates. Psychological review 119, 120.

14. Friedrich, J., and Lengyel, M. (2016). Goal-directed decision making with spiking neurons. Journal of Neuroscience 36, 1529–1546.

15. Zhang, Z., Cheng, Z., Lin, Z., Nie, C., and Yang, T. (2018). A neural network model for the orbitofrontal cortex and task space acquisition during reinforcement learning. PLOS Computational Biology 14, e1005925.

16. Rustichini, A., and Padoa-Schioppa, C. (2015). A neuro-computational model of economic decisions. Journal of neurophysiology 114, 1382–1398.

17. Song, H.F., Yang, G.R., and Wang, X. (2017). Reward-based training of recurrent neural networks for cognitive and value-based tasks. Elife 6, e21492.

18. Song, H.F., Yang, G.R., and Wang, X. (2016). Training excitatory-inhibitory recurrent neural networks for cognitive tasks: a simple and flexible framework. PLoS computational biology 12, e1004792.

19. Yang, G.R., and Wang, X. (2020). Artificial neural networks for neuroscientists: a primer. Neuron 107, 1048–1070.

20. Schulman, J., Wolski, F., Dhariwal, P., Radford, A., and Klimov, O. (2017). Proximal policy optimization algorithms. arXiv preprint 1707.06347.

21. Sutton, R., and Barto, A. (2018). Reinforcement learning: An introduction. MIT press.

22. Bongioanni, A., Folloni, D., Verhagen, L., Sallet, J., Klein-Flü gge, M.C., and Rushworth, M.F. (2021). Activation and disruption of a neural mechanism for novel choice in monkeys. Nature 591, 270–274.

23. Wang, X.J. (2002). Probabilistic decision making by slow reverberation in cortical circuits. Neuron 36, 955–968.

24. Wong, K.F., and Wang, X.J. (2006). A recurrent network mechanism of time integration in perceptual decisions. Journal of Neuroscience 26, 1314–1328.

25. Yang, G.R., Joglekar, M.R., Song, H., Newsome, W.T., and Wang, X. (2019). Task representations in neural networks trained to perform many cognitive tasks. Nature neuroscience 22, 297–306.

26. Driscoll, L.N., Shenoy, K., and Sussillo, D. (2024). Flexible multitask computation in recurrent networks utilizes shared dynamical motifs. Nature Neuroscience pp. 1–15.

27. Liu, Y., and Wang, X.J. (2024). Flexible gating between subspaces in a neural network model of internally guided task switching. Nature Communications 15, 6497.

28. Padoa-Schioppa, C. (2013). Neuronal origins of choice variability in economic decisions. Neuron 80, 1322–1336.

29. Roach, J.P., Churchland, A.K., and Engel, T.A. (2023). Choice selective inhibition drives stability and competition in decision circuits. Nature communications 14, 147.

30. Padoa-Schioppa, C. (2009). Range-adapting representation of economic value in the orbitofrontal cortex. Journal of Neuroscience 29, 14004–14014.

31. Kennerley, S.W., Behrens, T.E., and Wallis, J.D. (2011). Double dissociation of value computations in orbitofrontal and anterior cingulate neurons. Nature neuroscience 14, 1581–1589.

32. Dubreuil, A., Valente, A., Beiran, M., Mastrogiuseppe, F., and Ostojic, S. (2022). The role of population structure in computations through neural dynamics. Nature neuroscience 25, 783–794.

33. Fascianelli, V., Battista, A., Stefanini, F., Tsujimoto, S., Genovesio, A., and Fusi, S. (2024). Neural representational geometries reflect behavioral differences in monkeys and recurrent neural networks. Nature Communications 15, 6479.

34. Shi, W., Ballesta, S., and Padoa-Schioppa, C. (2022). Economic choices under simultaneous or sequential offers rely on the same neural circuit. Journal of Neuroscience 42, 33–43.

35. Pereira-Obilinovic, U., Hou, H., Svoboda, K., and Wang, X.J. (2024). Brain mechanism of foraging: Reward-dependent synaptic plasticity versus neural integration of values. Proceedings of the National Academy of Sciences 121, e2318521121.

36. Bengio, Y., Louradour, J., Collobert, R., and Weston, J. (2009). Curriculum learning. In Proceedings of the 26th annual international conference on machine learning. pp. 41–48.

37. Goudar, V., Peysakhovich, B., Freedman, D.J., Buffalo, E.A., and Wang, X.J. (2023). Schema formation in a neural population subspace underlies learning-to-learn in flexible sensorimotor problem-solving. Nature Neuroscience 26, 879–890.

38. Johnston, W.J., and Fusi, S. (2023). Abstract representations emerge naturally in neural networks trained to perform multiple tasks. Nature Communications 14, 1040.

39. Johnston, W.J., Fine, J.M., Yoo, S.B.M., Ebitz, R.B., and Hayden, B.Y. (2024). Semiorthogonal subspaces for value mediate a binding and generalization trade-off. Nature Neuroscience pp. 1–13.

40. O’Neill, M., and Schultz, W. (2010). Coding of reward risk by orbitofrontal neurons is mostly distinct from coding of reward value. Neuron 68, 789–800.

41. O’Neill, M., and Schultz, W. (2013). Risk prediction error coding in orbitofrontal neurons. Journal of Neuroscience 33, 15810–15814.

42. O’Neill, M., and Schultz, W. (2015). Economic risk coding by single neurons in the orbitofrontal cortex. Journal of Physiology-Paris 109, 70–77.

43. Ferrari-Toniolo, S., and Schultz, W. (2023). Reliable population code for subjective economic value from heterogeneous neuronal signals in primate orbitofrontal cortex. Neuron 111, 3683–3696.

44. Wang, X.J. (2022). Theory of the multiregional neocortex: large-scale neural dynamics and distributed cognition. Ann. Rev. Neurosci. 45, 533–560.

45. Schall, J.D., and Thompson, K.G. (1999). Neural selection and control of visually guided eye movements. Annual review of neuroscience 22, 241–259.

46. Cai, X., and Padoa-Schioppa, C. (2014). Contributions of orbitofrontal and lateral prefrontal cortices to economic choice and the good-to-action transformation. Neuron 81, 1140–1151.

47. Yim, M.Y., Cai, X., and Wang, X.J. (2019). Transforming the choice outcome to an action plan in monkey lateral prefrontal cortex: A neural circuit model. Neuron 103, 520–532.

48. Paszke, A., Gross, S., Massa, F., Lerer, A., Bradbury, J., Chanan, G., Killeen, T., Lin, Z., Gimelshein, N., Antiga, L. et al. (2019). Pytorch: An imperative style, high-performance deep learning library. Advances in neural information processing systems 32.

49. Huang, S., Dossa, R.F.J., Ye, C., Braga, J., Chakraborty, D., Mehta, K., and Araújo, J.G. (2022). Cleanrl: High-quality single-file implementations of deep reinforcement learning algorithms. Journal of Machine Learning Research 23, 1–18. URL: http://jmlr.org/papers/v23/21-1342.html.

50. Brockman, G., Cheung, V., Pettersson, L., Schneider, J., Schulman, J., Tang, J., and Zaremba, W. (2016). Openai gym. 1606.01540.

51. Harris, C.R., Millman, K.J., van der Walt, S.J., Gommers, R., Virtanen, P., Cournapeau, D., Wieser, E., Taylor, J., Berg, S., Smith, N.J., Kern, R., Picus, M., Hoyer, S., van Kerkwijk, M.H., Brett, M., Haldane, A., del Río, J.F., Wiebe, M., Peterson, P., Gérard-Marchant, P., Sheppard, K., Reddy, T., Weckesser, W., Abbasi, H., Gohlke, C., and Oliphant, T.E. (2020). Array programming with NumPy. Nature 585, 357–362. URL: https://doi.org/10.1038/s41586-020-2649-2. xdoi: 10.1038/s41586-020-2649-2.

52. Pedregosa, F., Varoquaux, G., Gramfort, A., Michel, V., Thirion, B., Grisel, O., Blondel, M., Prettenhofer, P., Weiss, R., Dubourg, V., Vanderplas, J., Passos, A., Cournapeau, D., Brucher, M., Perrot, M., and Duchesnay, E. (2011). Scikit-learn: Machine learning in Python. Journal of Machine Learning Research 12, 2825–2830.

53. Markram, H., Toledo-Rodriguez, M., Wang, Y., Gupta, A., Silberberg, G., and Wu, C. (2004). Interneurons of the neocortical inhibitory system. Nature reviews neuroscience 5, 793–807.

54. Douglas, R.J., and Martin, K.A. (2007). Recurrent neuronal circuits in the neocortex. Current biology 17, R496–R500.

55. Schultz, W., Dayan, P., and Montague, P.R. (1997). A neural substrate of prediction and reward. Science 275, 1593–1599.

56. Glimcher, P.W. (2011). Understanding dopamine and reinforcement learning: the dopamine reward prediction error hypothesis. Proceedings of the National Academy of Sciences 108, 15647–15654.

57. Kingma, D., and Ba, J. (2014). Adam: A method for stochastic optimization. arXiv preprint 1412.6980.

