## Supplemental Material for "A neural circuit framework for economic choice: from building blocks of valuation to compositionality in multitasking"

November 25, 2025

### Supplementary Figures

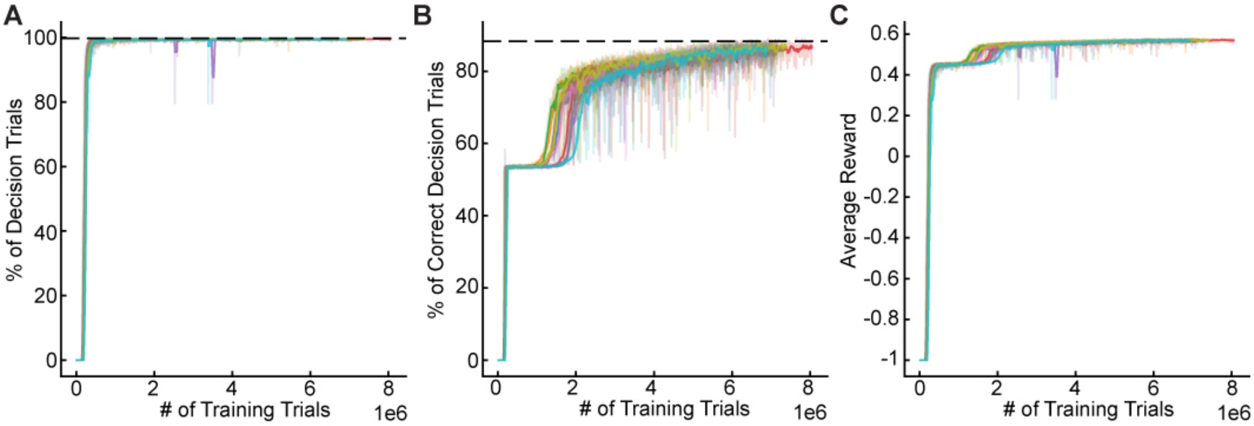


**Figure S 1. Learning curves for networks trained on all tasks.** (A) Percentage of decision trials (e.g., trials completed without fixation breaks) on the test set as a function of training trials. The black dashed line indicates the learning criterion of 99% decision trials. (B) Percentage of correct choices among decision trials on the test set as a function of training trials. The black dashed line indicates the learning criterion of 90% correct decisions. (C) Average reward on the test set as a function of training trials. In all panels, percentages and average rewards are computed on test trials. Colored lines represent different networks with random initializations; solid lines are smoothed versions. The learning curves show that networks typically progressed through three distinct phases: first, learning to maintain fixation; second, learning to select an action randomly; and finally, learning the value-based policy of selecting the highest-value offer. Training stops once both criteria are met.

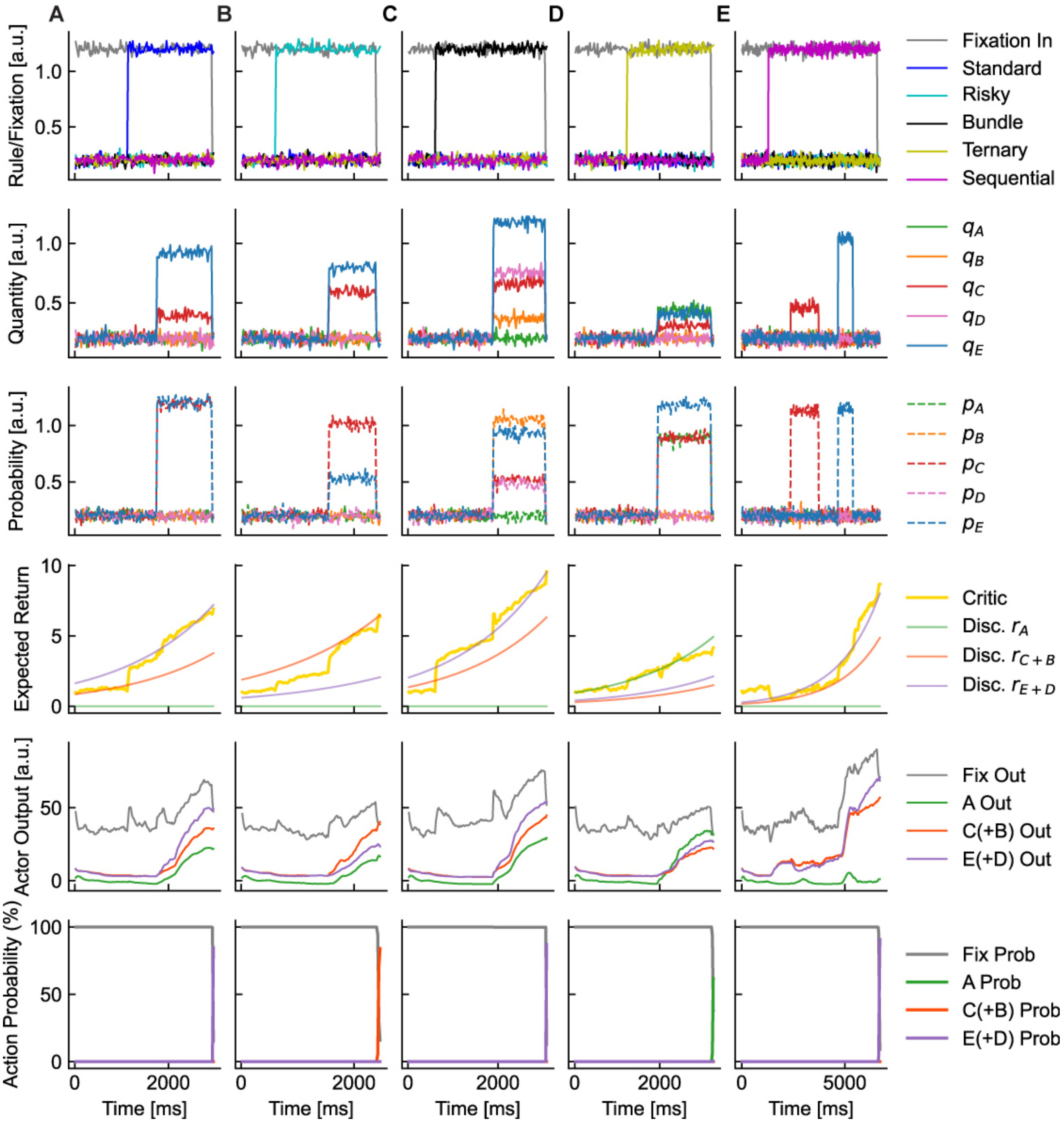


**Figure S 2. Example trials from a trained network on the five economic choice tasks.** Each column (A-E) corresponds to a sample trial from one of the five tasks: (A) Standard, (B) Risky, (C) Bundles, (D) Ternary, and (E) Sequential. Within each trial, the rows display the temporal dynamics of different signals. The first row shows fixation and rule cue inputs. The second and third rows show the quantities and probabilities of the offered goods, respectively. The fourth row shows the value function output (critic) and potential discounted rewards. The fifth and sixth rows illustrate the policy outputs (logits) and the resulting softmax probabilities for the network's possible actions. Action labels like "C(+B) Out" refer to the action of choosing Good C or the bundle containing it, depending on the task. In each trial, the network correctly selects the highest-value offer. The policy outputs are dominated by fixation during the offer period, but the network's forthcoming choice can often be inferred from the rising probability of the chosen action before the response is executed. The trials have varying epoch durations, reflecting the variability in the task design.

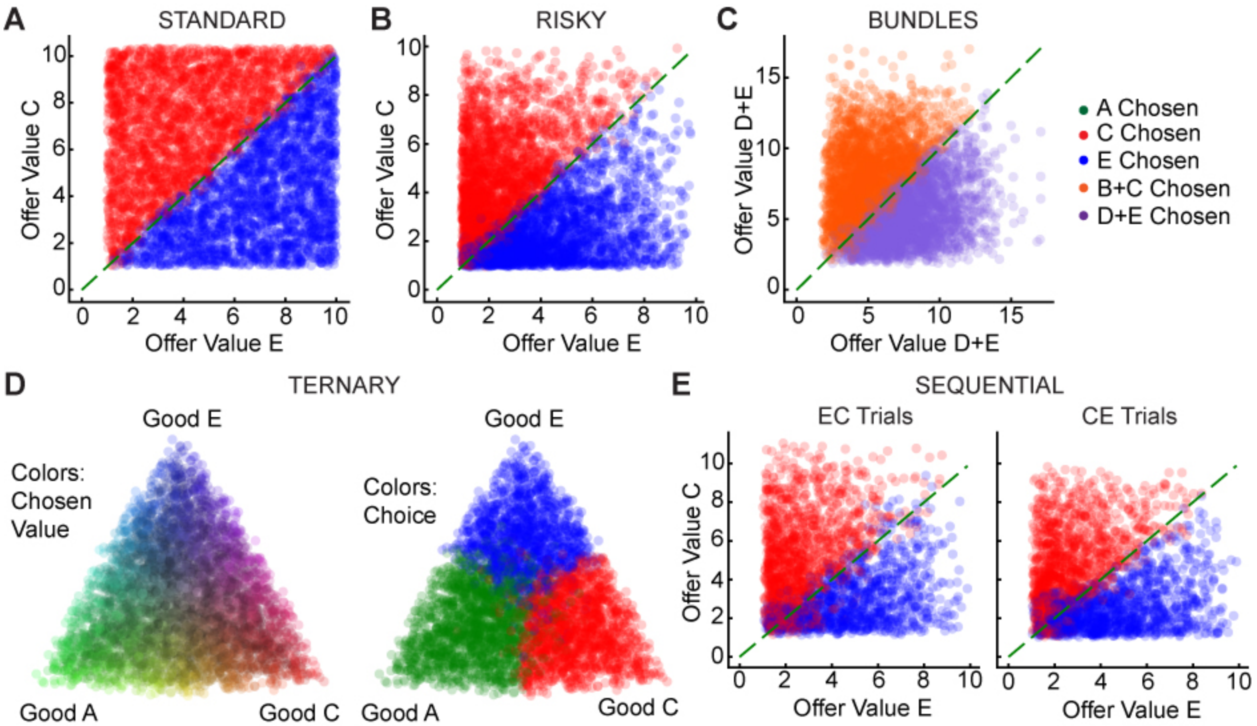


**Figure S 3. Choice patterns of a network trained on all economic choice tasks.** Each panel displays the network's choices in the computed offer value space for a different task, demonstrating a consistent value-based strategy. Each point is a single trial; the dashed green line is the identity line where both offers have equal value. (A) *Standard task:* Choices between goods C (red) and E (blue) are cleanly separated by the offer value identity line. (B) *Risky task:* As in the main figure, choices are linearly separable in offer value space. (C) *Bundles task:* Choices between a bundle of goods B+C (orange) and a bundle of goods D+E (purple) are consistently determined by the total summed value of each bundle. (D) *Ternary task:* Choices among three goods (A, C, E) are visualized on a simplex. Each trial's position is determined by the relative offer values of the three goods. Left: Trials are colored by the mixture of offer values (A=green, C=red, E=blue), with the dominant color indicating the highest-value good. Right: The same trials are colored by the network's choice. The close correspondence between the colors in the two plots shows that the network reliably selects the good with the highest offer value. (E) *Sequential task:* Choices between sequentially presented goods C and E. The network accurately compares the current offer with the one held in memory, irrespective of presentation order (EC trials: E first; CE trials: C first).

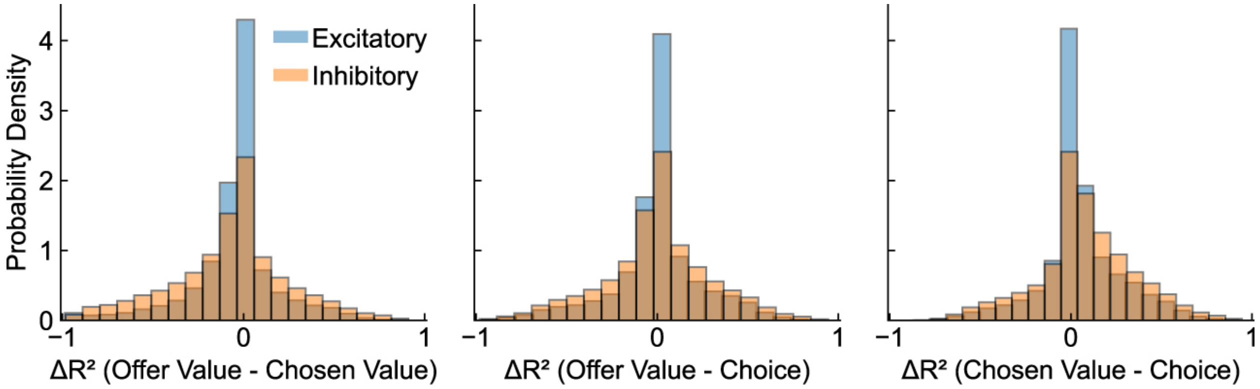


**Figure S 4. Widespread mixed selectivity in the general population of task-modulated neurons.** Histograms show the difference in $R^2$ values ($\Delta R^2$) between pairs of decision variables, analogous to Figure 3E in the main text. However, this analysis includes all neurons that were significantly modulated by at least one variable ($p < 0.05$ and a lenient $R^2 \geq 0.005$). The resulting unimodal distributions, sharply peaked at zero, indicate that most task-related neurons exhibit mixed selectivity, responding to combinations of variables rather than encoding them categorically. This suggests the network employs a dual coding strategy, with a large population of mixed-selectivity neurons and a smaller, specialized subpopulation of categorical neurons (Figure 3E).

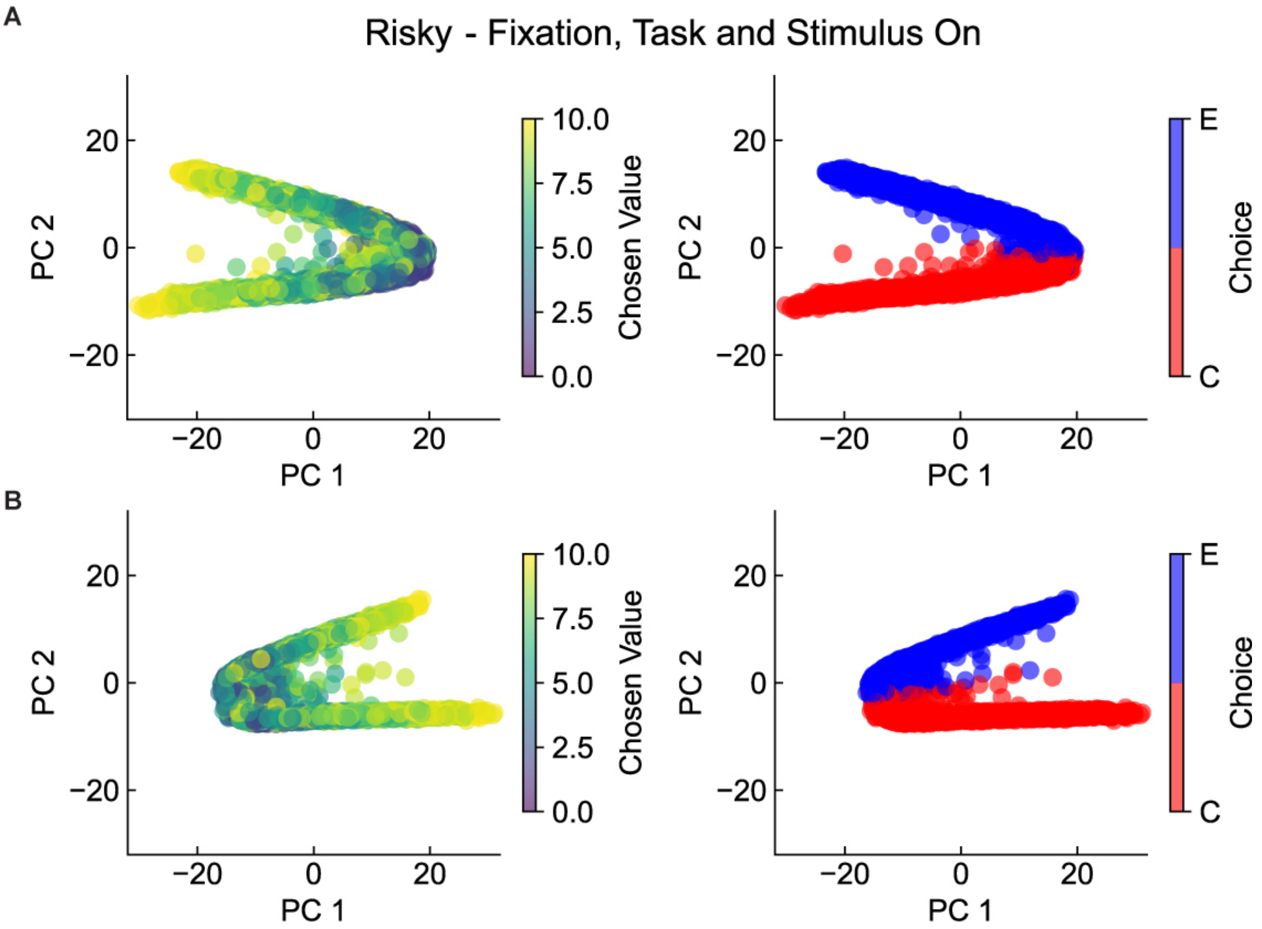


**Figure S 5. Population dynamics were qualitatively consistent across independently trained networks.** (A-B) Snapshots of population activity from two different, representative networks (other than the one shown in Figure 4) during the risky task. Although the exact orientation of the principal components could vary due to rotational symmetries, all networks learned a similar low-dimensional solution where the cloud of population states separated based on the upcoming choice. This demonstrated that the emergent population dynamics represented a robust and convergent solution.

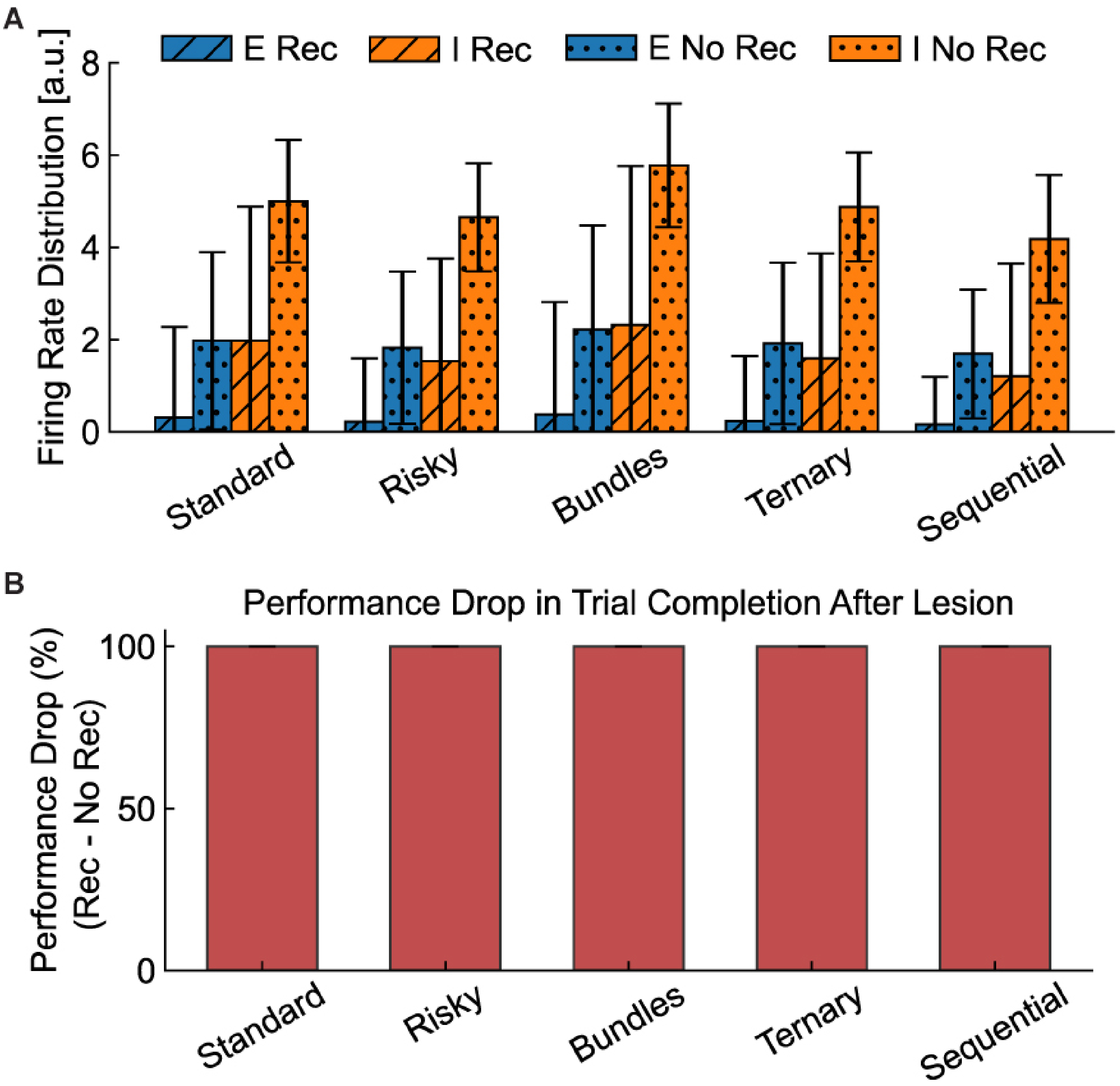


**Figure S 6. Lesioning recurrent connections dysregulated neural activity and completely abolished choice behavior.** (A) Distribution of firing rates for excitatory and inhibitory neurons in intact (Rec) versus lesioned (No Rec) networks. Removing recurrent connections led to an increase in firing rates due to the loss of recurrent inhibition and competition. Error bars represent standard deviation. (B) Behavioral performance after lesioning all recurrent connections. The network failed to make a choice in 100% of trials, resulting in a complete performance drop where all trials were aborted. This demonstrated the critical and profound behavioral role of the recurrent circuitry for executing a choice.

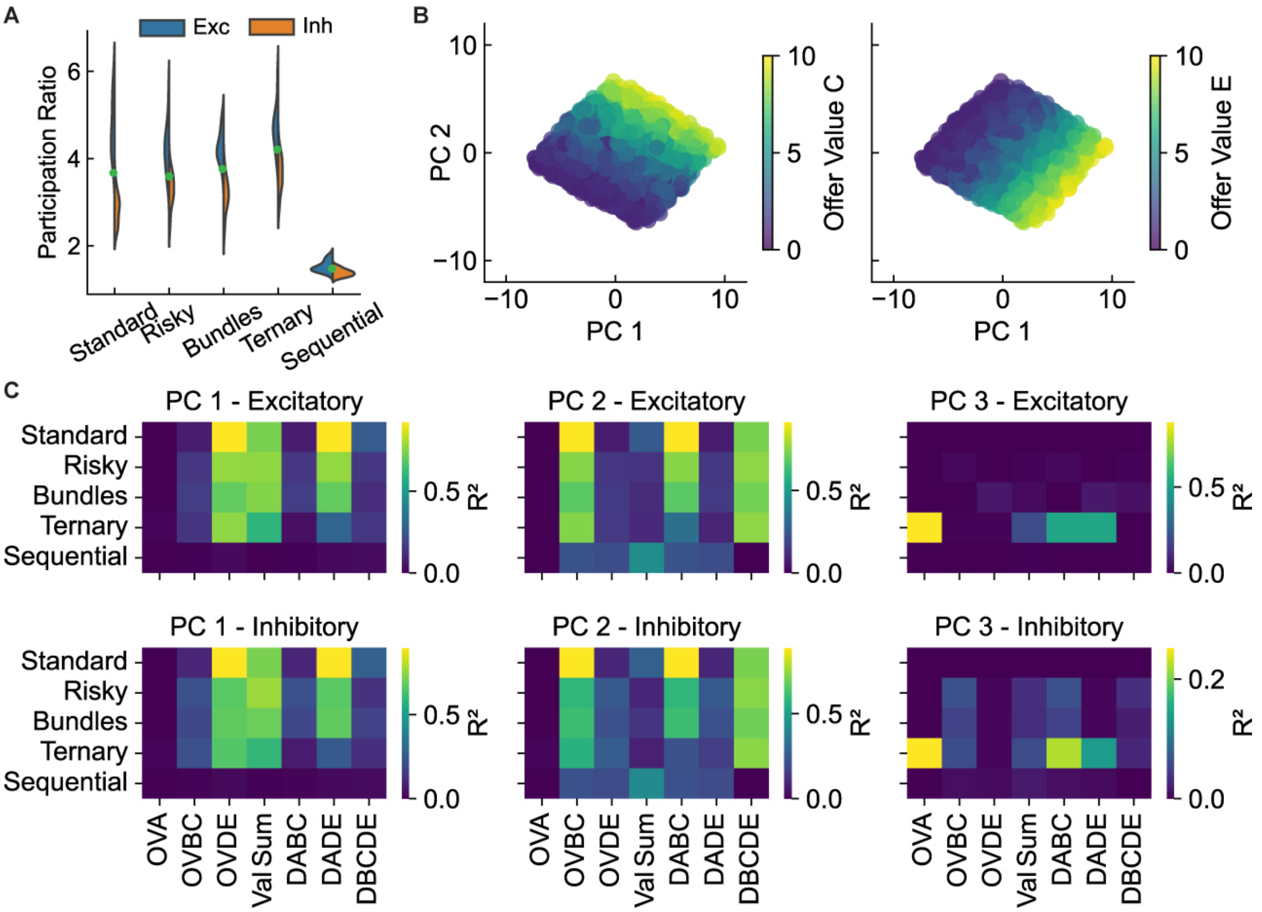


**Figure S 7. Lesioned networks retained feedforward value signals but lost the capacity for value comparison.** (A) Neural dimensionality in lesioned networks. Compared to the intact networks (Fig. 4A), the dimensionality for the sequential task was substantially lower, consistent with a loss of working memory without recurrent connections. Conversely, dimensionality for the other tasks was higher, suggesting that the winner-take-all dynamics in the intact circuit served to compress information about the unchosen offers. (B) Snapshots of population activity in a representative lesioned network. The population states were still organized by offer value (left: colored by Offer Value C; right: colored by Offer Value E), but crucially, they failed to separate based on the eventual (but unmade) choice. This visually demonstrated the loss of the choice representation. (C) Summary of population analyses across all lesioned networks. The set of regressors was critically different from the intact network analysis because the lesioned network could not make a choice, and thus post-decision variables were absent. Regressions were performed against input-related variables: offer values ($OVA$, $OVBC$, $OVDE$), their sum ($Val\ Sum$), and their differences ($D...$). For example, $DBCDE$ represents the value Difference between bundle B+C and bundle D+E. The heatmaps show that the population still encoded input variables (offer values, sum, and differences) but had completely lost the representation of chosen value and choice (which were not included as regressors because they could not be computed). This causally demonstrated that recurrence was necessary for value comparison, not value representation itself.

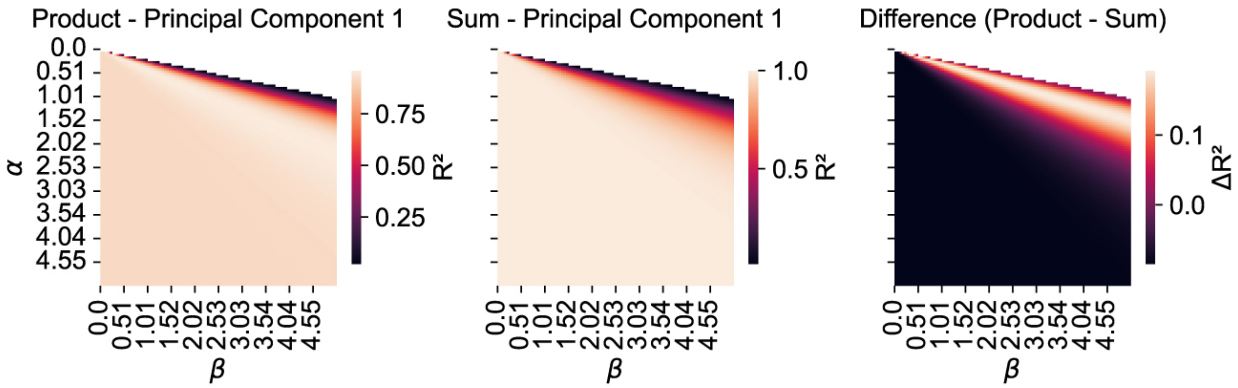


**Figure S 8. Parameter search for the toy multiplication model.** This figure identifies the optimal parameters for the simplified feedforward model to approximate a multiplicative function, justifying the parameters used in Fig. 5C-D. We systematically varied the scaling parameters for the input weights ($\alpha$) and biases ($\beta$) of the model's hidden units. For each parameter combination, we evaluated how well the resulting population activity could be linearly decoded to represent either the product or the sum of the inputs. Left panel: The heatmap shows the coefficient of determination ($R^2$) for a linear regression of the first principal component of the hidden units' activity against the *product* of the inputs. Center panel: The same analysis for the *sum* of the inputs. Right panel: The difference between the two $R^2$ values ($\Delta R^2 = R^2_{\text{product}} - R^2_{\text{sum}}$). The bright region of the right panel highlights the parameter regime where the model's internal representation was best explained by multiplication, significantly outperforming an additive model. The parameters used for the toy model simulations in Figure 5 were selected from this optimal region.

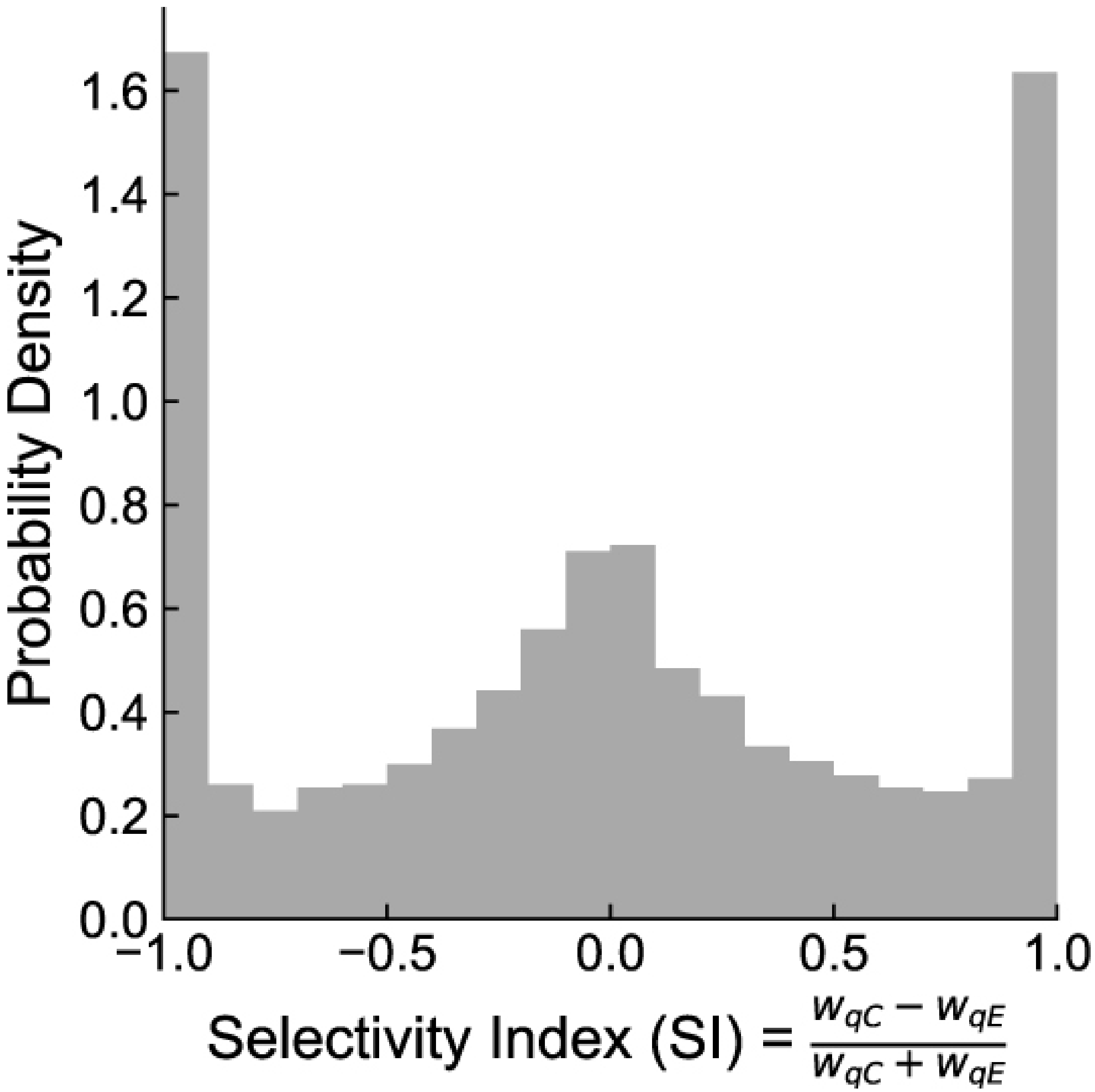


**Figure S 9. Direct evidence for a dual architecture of specialized and mixed-selective input pathways.** This histogram reveals the underlying circuit architecture for feedforward value processing by analyzing the input synaptic weights from the two competing offers (C and E). We calculated a Selectivity Index (SI) for each recurrent neuron, defined as $SI = (w_{qC} - w_{qE})/(w_{qC} + w_{qE})$. The distribution of SI values across the population reveals three distinct features: two sharp peaks at the extremes (SI = $-1$ and $+1$), corresponding to two populations of neurons that are highly specialized for a single offer, and a third, large, broad unimodal distribution centered near zero. This central peak provides direct evidence for a substantial population of mixed-selective neurons that integrated information from both offers. This dual architecture, combining both specialized and integrated pathways, provided the necessary anatomical substrate for the network to simultaneously represent individual offer values and compute their sum, which causally generated the geometric rotation to a value sum/difference representation (Fig. 5D). The histogram shows the average probability density across the 50 networks trained exclusively on the risky task.

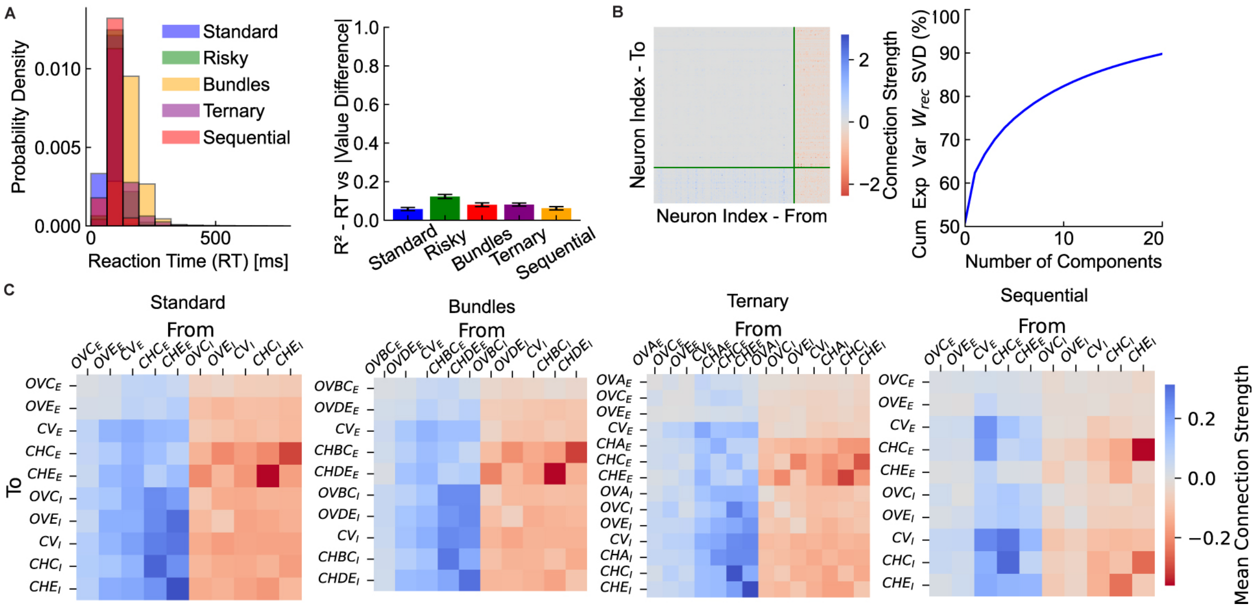


**Figure S 10. Additional analyses of choice mechanisms and connectivity.** (A) Left: Distributions of reaction times (RTs) showed similar profiles across all tasks. Right: Linear regression of RT vs. the absolute value difference between offers showed no significant relationship, consistent with an early commitment to a choice. (B) Left: An example of a full 256 × 256 recurrent connectivity matrix from a single representative trained network. Right: Cumulative explained variance from Singular Value Decomposition (SVD) of the weight matrix, averaged across all 20 multitasking networks, revealed a consistent low-rank structure. This indicates that the complex full matrix can be well approximated by a few dominant components, which correspond to the powerful, stereotyped connectivity motifs that govern network dynamics. (C) Mean reduced connectivity matrices for the standard, bundles, ternary, and sequential tasks, averaged across all 20 networks. This analysis showed that the same core CRI motif emerged as a general solution across different choice contexts.

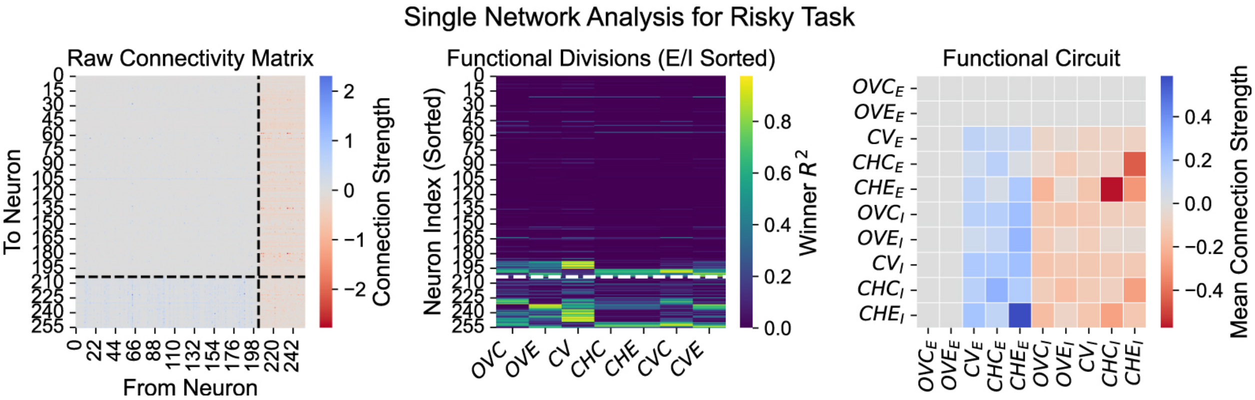


**Figure S 11. Derivation of the reduced connectivity matrix via many-to-one functional abstraction.** This schematic illustrates the three-step pipeline used to derive the interpretable circuit diagrams from the full recurrent weight matrix for a single example network. *Left (Raw Connectivity Matrix):* The full $256 \times 256$ weight matrix, with connections from excitatory (neurons $0 - 203$) and to inhibitory (neurons $204 - 255$) neurons segregated. *Center (Functional Divisions):* Each neuron (y-axis, sorted by E/I type with dotted white line showing boundary) was classified based on which decision variable (x-axis) its activity best explained (color indicates $R^2$ value). *Right (Functional Circuit):* The effective connection strength between each functional pool was calculated by averaging all individual synaptic weights between them. This single-network matrix, when averaged with others, produced the final reduced matrix in Fig. 6C.

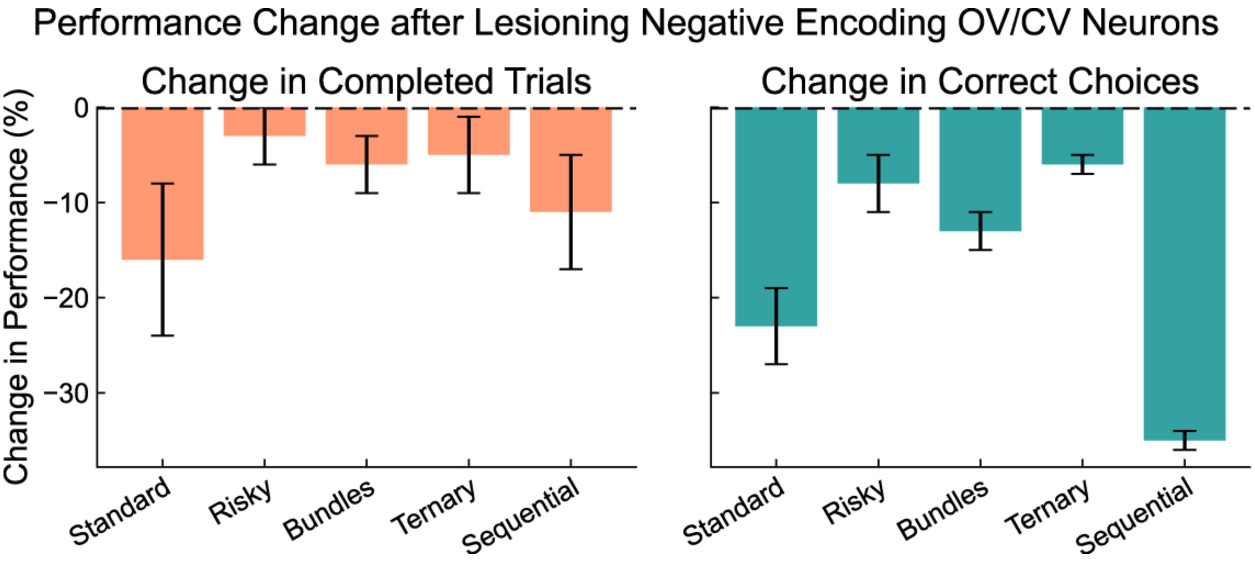


**Figure S 12. Causal role of negatively tuned neurons in robust decision-making.** To test the functional importance of negatively tuned neurons, we performed a targeted lesion analysis. We computationally silenced all neurons with a significant negative encoding of either offer value or chosen value and evaluated performance on all five tasks. The bar plots show the percentage change in performance for (Left) Completed Trials and (Right) Choice Accuracy (on completed trials), relative to the intact network. The lesion significantly impaired performance across all tasks. This demonstrated that negatively tuned neurons were computationally essential for robust decision-making.

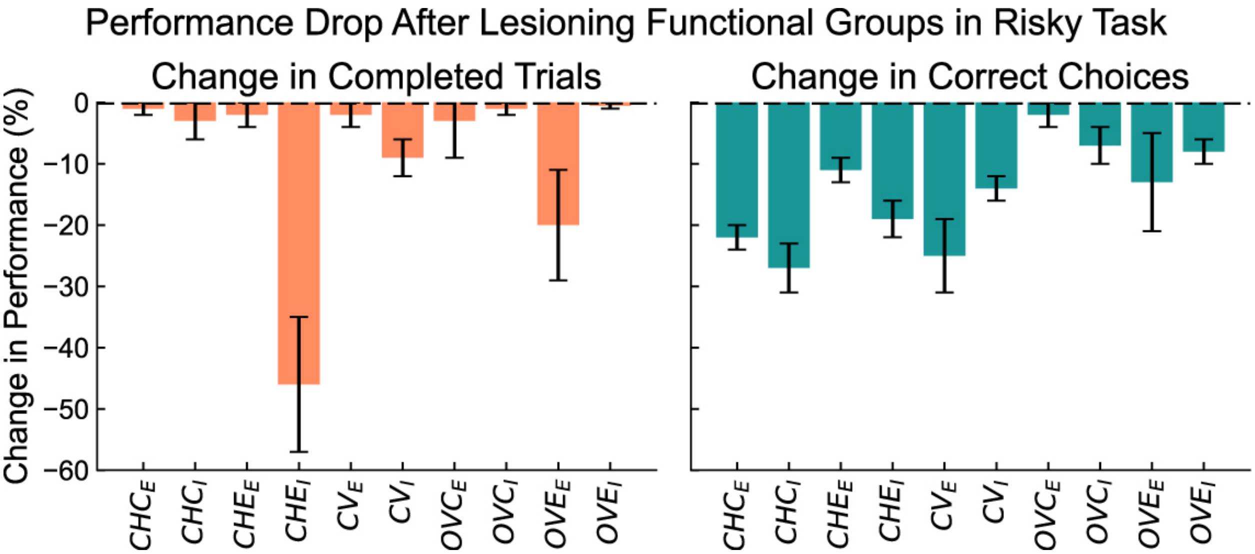


**Figure S 13. Causal validation of decision-critical functional cell groups.** To identify the most critical neural populations for value comparison, we performed a systematic functional ablation analysis on the 20 multitask networks, tested on the risky task. Each functional pool of neurons, classified by its primary selectivity, was computationally silenced. The bar plots show the resulting percentage change in (Left) Completed Trials and (Right) Choice Accuracy (on completed trials). The results highlighted the distributed nature of the circuit's function. While lesioning the choice-selective inhibitory population '$CHE_I$' had a uniquely large effect on trial completion, deficits in choice accuracy were severe after lesioning any of the choice-selective populations (both excitatory '$CHC_E$', '$CHE_E$' and inhibitory '$CHC_I$', '$CHE_I$') as well as the chosen-value excitatory ('$CV_E$') population. Notably, the impact of silencing choice-selective inhibitory pools on accuracy was as large as silencing their excitatory counterparts, causally identifying the full E/I choice-selective circuit as the core pillar of the decision-making process. Acronyms: OVC/E (Offer Value C/E), CV (Chosen Value), CHC/E (Choice C/E). Subscripts $_E$ and $_I$ denote excitatory and inhibitory populations.

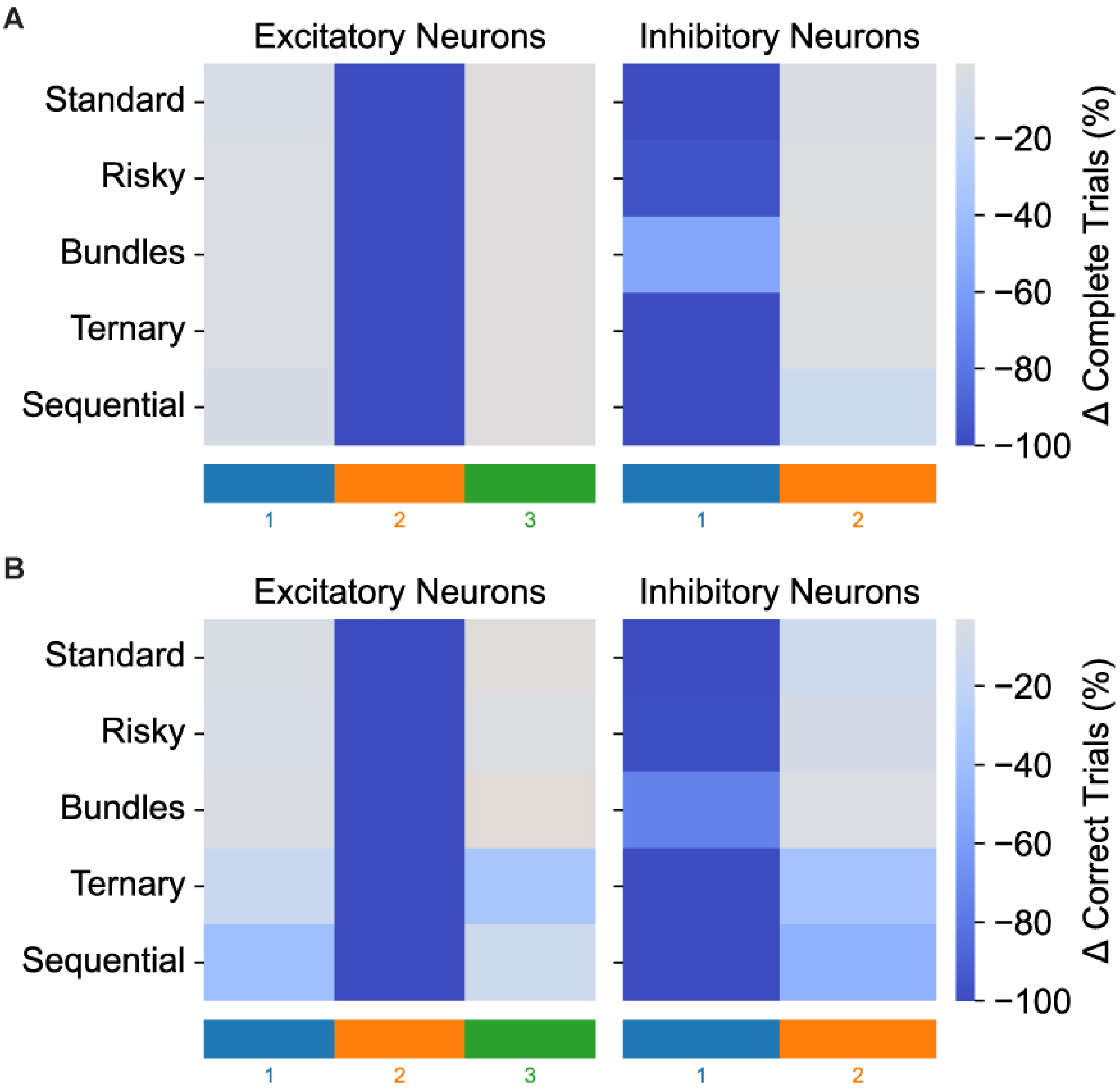


**Figure S 14. Task-specific deficits following targeted lesions of functionally specialized neuronal clusters.** This figure provided causal evidence for the functional roles of the neuronal clusters identified in Figure 7C. The heatmaps show the percentage drop in performance ((A): Trial Completion Rate; (B): Accuracy on Completed Trials) after computationally silencing all neurons within a specific cluster. Each column block corresponds to the lesion of a single cluster. Lesioning the shared cluster caused a global, catastrophic deficit across all five tasks. In contrast, lesioning the ternary-specialized cluster selectively impaired performance primarily on the ternary task. Lesioning the sequential-specialized cluster caused a performance drop almost exclusively in the sequential task. This systematic analysis provided direct causal evidence for a functionally specialized, compositional architecture.

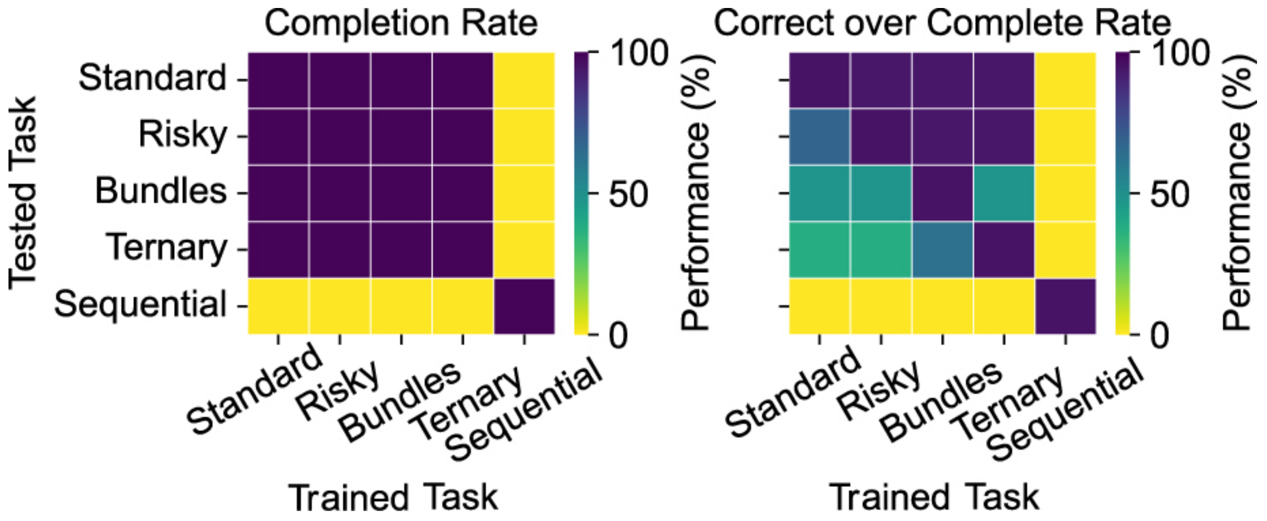


**Figure S 15. Zero-shot generalization performance of single-task trained networks.** The matrices show the mean performance of networks trained exclusively on one task (x-axis) when tested on all other tasks (y-axis), with 10 networks per trained task group. The left panel shows the Trial Completion Rate, while the right panel shows the Accuracy on successfully completed trials. High performance on the diagonal (dark purple, $\sim$ 100%) confirmed successful training on the source task. The off-diagonal elements revealed strongly asymmetric generalization. For instance, networks trained on a complex task like Ternary could perform simpler, subsumed tasks (dark colors in the Ternary column). Conversely, networks failed on tasks requiring new computational modules, such as a working memory component (bright yellow, indicating $\sim$ 0% completion rate, for most networks tested on the Sequential task). This causally demonstrated that skills are learned as distinct, compositional modules and that multitasking is necessary to build a flexible, multi-skilled circuit.

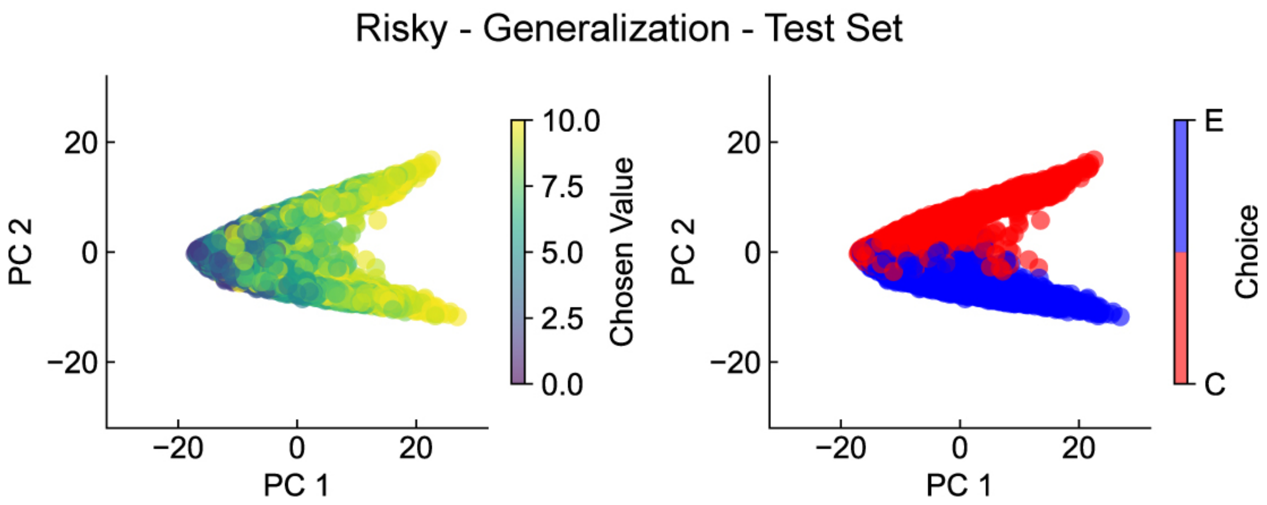


**Figure S 16. Neural dynamics of decision-making generalize to novel offers.** This figure shows that the low-dimensional geometric solution for decision-making is preserved when networks generalize to novel stimuli they have never encountered before. Principal component analysis was performed on the population activity of a representative constrained-trained network during the generalization test phase. Each point represents the population state of a single trial projected onto the first two principal components. Left panel: When colored by the chosen value, the population states show a clear gradient along PC1, indicating that this axis encodes the value of the selected offer. Right panel: When the same states are colored by the network's choice (C in red, E in blue), the data cloud separates into two distinct clusters along PC2. This demonstrates that the network applies the same geometric solution observed in standard-trained networks (Fig. 4B): it forms a decision by driving its neural state into one of two choice-specific manifolds, with the final position along the manifold determined by the chosen value. This provides a robust neural basis for the network's ability to generalize its decision-making capability.

### Supplementary Videos

**Supplementary Video S1. Dynamic visualization of decision formation in an intact network.** The video shows the temporal evolution of the population state cloud (each point is a single trial) during the risky task, projected onto the first two principal components. The four panels show the same dynamics, colored by different decision variables. Following stimulus onset, the population trajectories evolve from a common starting point and dynamically separate into distinct red and blue clusters in the 'Choice' panel (bottom right), vividly illustrating the formation of the decision in the neural state space.

**Supplementary Video S2. Dynamic visualization of failed decision formation in a lesioned network.** The video shows the evolution of the population state in a network with all recurrent connections removed. Each point is a single trial's state. While the internal structure of the cloud is graded by the individual offer values, it fails to separate into distinct choice clusters over time. This visually demonstrates the network's inability to perform value comparison and form a decision without the competitive dynamics mediated by the recurrent circuitry.
